# Mycobacterium tuberculosis reactivates HIV via exosomes mediated resetting of cellular redox potential and bioenergetics

**DOI:** 10.1101/629048

**Authors:** Priyanka Tyagi, Virender Kumar Pal, Sandhya Srinivasan, Amit Singh

## Abstract

The synergy between *Mycobacterium tuberculosis* (*Mtb*) and HIV-1 interferes with therapy and facilitates pathogenesis of both human pathogens. Fundamental mechanisms by which *Mtb* exacerbates HIV-1 are not clear. Here, we show that exosomes secreted by macrophages infected with *Mtb*, including drug-resistant clinical strains, reactivate HIV-1 by inducing oxidative stress. Mechanistically, *Mtb*-specific exosomes realign mitochondrial and non-mitochondrial oxygen consumption rate (OCR) and modulates the expression of genes mediating oxidative stress response, inflammation, and HIV-1 transactivation. Proteomics revealed the enrichment of several host factors (*e.g.,* HIF-1α, galectins, Hsp90) known to promote HIV-1 reactivation in the *Mtb*-specific exosomes. Treatment with a known antioxidant, N-acetyl cysteine, or with the inhibitors of host factors galectins and Hsp90 attenuated HIV-1 reactivation by *Mtb*-specific exosomes. Our findings uncovered new paradigms for understanding the redox and bioenergetics basis of HIV-TB co-infection, which will enable the design of effective therapeutic strategies.

## Introduction

Tuberculosis (TB) and Human Immunodeficiency Virus/Acquired Immunodeficiency Syndrome (HIV/AIDS) jointly represent the major burden of infectious diseases in humans worldwide. HIV-1/*Mtb* co-infected patients exhibit rapid progression to AIDS and shorter survival kinetics [1]. Moreover, the risk of acquiring active TB infection increases from 10% in a lifetime to 10% per year in the case of HIV-1 infected patients [2]. In 2015 alone, WHO estimated that almost 11% of 10.4 million TB patients are also infected with HIV-1, and one in every three deaths between HIV-1 infected individuals was due to TB (http://www.who.int/hiv/topics/tb/about_tb/en/). Epidemiological studies clearly indicate that both these human pathogens interact to accelerate disease severity and deaths [2].

Since infection with HIV-1 significantly increases the risk of TB reactivation in individuals latently infected with *Mtb* [3], a great majority of studies were focused on revealing how virus disorganizes TB granuloma [4], impaired phagosomal killing [5], and alters T-cell based immunity to exacerbates *Mtb* pathogenesis [6]. In contrast, whether *Mtb* influences exit of HIV-1 from latency and it’s reentry into a productive life cycle remain poorly studied. Because HIV-1 can persist in a latent state for decades and then reactivates to cause immunodeficiency, our particular interest is to understand the mechanism, if any, underlying *Mtb* induced reactivation of HIV-1 from latency. Growing body of evidences suggest that infection with *Mtb* or its component(s) (lipids and secretory proteins) promotes HIV-1 replication by regulating processes such as inflammation, MHCII processing, Toll-Like Receptors (TLRs) signaling, CXCR4/CCR5 expression, proinflammatory cytokines/chemokines production, and activating transcriptional regulators (NF-kB, NFAT) of the long-terminal repeats (LTRs) of HIV [7-13]. Contradictory evidences showing inhibition of HIV-1 replication by *Mtb* complicates our understanding of how two human pathogens interact at the molecular level [14, 15]. Despite this, research specifically addressing how *Mtb* modulates HIV latency and reactivation is, however, quite scarce. In this context, reactive oxygen species (ROS) and modulation of central metabolism are considered to be one of the main mechanisms regulating HIV-1 replication, immune dysfunction, and accelerated progression to AIDS [16]. Deeper studies in this direction revealed an important role for a major cellular antioxidant, glutathione (GSH) [17]. Low GSH levels in HIV patients have been shown to induce provirus transcription by activation of NFκB, apoptosis of CD4 T cells, and depletion of CD4 T cells [18]. Consequently, replenishment of GSH is considered as a potential supplement to highly active antiretroviral therapy (HAART) [19]. Recently, we have reported subtle changes in the intracellular and subcellular redox potential of GSH (*E_GSH_*) modulates HIV-1 replication cycle [20]. We discovered that oxidative stress caused by a marginal increase in intracellular *E_GSH_*(25 millivolts [mV]) is sufficient to trigger HIV-1 reactivation from latently infected cells, raising the potential of targeting HIV-1 latency by the modulators of cellular GSH homeostasis [20]. Interestingly, markers of oxidative stress such as ROS/RNS and lipid peroxidation have been shown to be elevated in active TB patients [21]. Specifically, serum/cellular GSH was either depleted or oxidized in human TB patients and in the lungs of guinea pigs infected with *Mtb* [21, 22]. Treatment with GSH precursor, N-acetyl cysteine (NAC), reversed oxidative stress to reduce bacterial survival and tissue damage in guinea pigs [22]. Additionally, *Mtb* infection has recently been shown to influence carbon flux through glycolysis and TCA cycle in infected macrophages [23]. This, along with the recognized role of GSH homeostasis and glycolysis in HIV infection, indicates that the two pathogens might synergies via affecting redox signaling and metabolic phenotypes of the host. We explored this connection and investigated if *Mtb* induces variation in the intracellular *E_GSH_*and bioenergetics to induce HIV-1 reactivation program. We anticipate that such a study has the potential to overcome many of the deficiencies in our understanding of the metabolic basis of HIV-TB co-infection and may enable high throughput screen to identify small molecule modulators of redox/central metabolism to potentiate the intervention strategies against HIV-TB co-infection.

In this study, we showed that macrophages infected with virulent *Mtb,* including multi-drug resistant (MDR) and extensively drug-resistant (XDR) clinical strains, release exosomes to induce substantial oxidative stress in neighboring non-infected macrophages (bystander). Taking cues from these findings, we showed that *Mtb* exploits the exosome-based mechanism to communicate and reactivate latent HIV-1 from monocytic (U1) and lymphocytic (J-Lat) cells. Mechanistically, *Mtb* specific exosomes alter gene expression, redox metabolism, and bioenergetics of latent cells to promote HIV-1 reactivation. Proteomic analysis of exosomes indicates that *Mtb* infection transports several proteins associated with host cellular pathways known to reactivate HIV-1 by perturbing redox metabolism, inflammation, and immune response.

## Results

### *Mtb* infection induces oxidative stress in bystander macrophages

We exploited a non-invasive biosensor (Grx1-roGFP2) of GSH redox potential (*E_GSH_*) [24] to measure dynamic changes in the redox physiology of human macrophages (U937) upon infection with a virulent strain of *Mtb* (H37Rv). GSH is the most abundant low molecular weight thiol produced by mammalian cells, therefore, *E_GSH_* measurement provides a reliable and sensitive indicator of the cytoplasmic redox state of macrophages [20, 24]. The biosensor shows an increase in the fluorescence excitation ratio at 405/488 nm upon oxidative stress, whereas a ratiometric decrease is associated with reductive stress (Fig. 1A). The ratiometric changes can be easily fitted into the modified Nernst equation to precisely calculate *E_GSH_* [24].

**Fig. 1.**
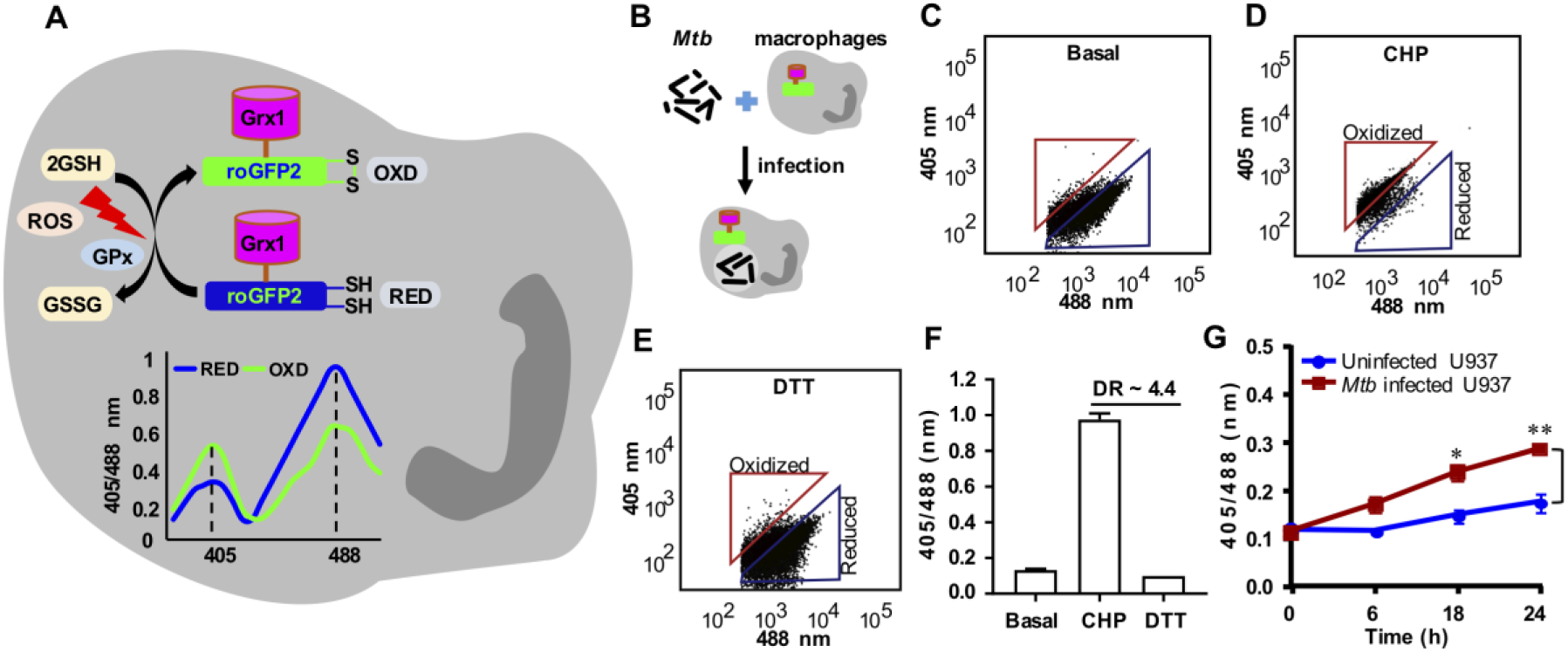
*Mycobacterium tuberculosis (Mtb)* induces oxidative shift in *E_GSH_*of U937 macrophages (Mφ). **(A)** Schematic representation of Grx1-roGFP2 oxidation and reduction in response to ROS inside a mammalian cell stably expressing the biosensor. GPx denotes GSH-dependent glutathione peroxidase. The graph represents the 405/488 nm ratios change upon oxidation or reduction of Grx1-roGFP2 in response to oxidative or reductive stress. Oxidative stress increases fluorescence at 405 nm excitation and decreases fluorescence at 488 nm at a constant emission of 510 nm, whereas an opposite biosensor response is induced by reductive stress. **(B)** PMA differentiated U937 Mφ stably expressing Grx1-roGFP2 in the cytosol were infected with *Mtb* H37Rv at a multiplicity of infection (MOI) 10. At the indicated time points, ratiometric sensor response was measured by exciting the sensor at 405 and 488 nm lasers and at a constant emission of 510 nm using flow cytometer. Dot plot spectra showing the ratiometric shift in the biosensor response in case of **(C)** untreated U937 (basal) and upon treatment of with **(D)** an oxidant cumene hydroperoxide (CHP, 0.5 mM) and **(E)** a reductant dithiothreitol (DTT, 40 mM). **(F)** Dynamic range (DR) of the biosensor in U937 cells based on the complete oxidation and reduction by CHP and DTT, respectively. **(G)** Ratiometric biosensor response over time in case of uninfected and *Mtb* infected U937 macrophages. Error bars represent standard deviations from the mean. *p<0.05; **p<0.01 (two-way ANOVA). Data are representative of at least three independent experiments performed in triplicate.

U937 monocytes expressing cytosolic Grx1-roGFP2 (U937/Grx1-roGFP2) were differentiated to macrophages using PMA and infected with *Mtb* H37Rv (Fig. 1B). At various time-points post-infection, 405/488 ratios were measured by flow cytometry to calculate intracellular *E_GSH_* as described [20]. We first confirmed the response of biosensor to a well-known oxidant cumene hydroperoxide (CHP) and a cell-permeable thiol-reductant dithiothreitol (DTT). As expected, the treatment of U937/Grx1-roGFP2 with CHP increases 405/488 ratio, which corresponds to *E_GSH_*of −240 mV and treatment with a cell-permeable thiol-reductant dithiothreitol (DTT) decreases 405/488 ratio, which corresponds to *E_GSH_* of −320 mV. (Fig. 1C-F). Next, we examined the biosensor response upon infection with *Mtb* H37Rv. Uninfected U937/Grx1-roGFP2 cells exhibit highly reduced cytoplasm (405/488 ratio ∼ 0.1-0.15 over-time; *E_GSH_* = −301 ± 2 mV) (Fig. 1G). In contrast, *Mtb* infection gradually increases the biosensor oxidation ratio over time, which results in an ∼ +20 mV shift in *E_GSH_*(−282 ± 2 mV) at 24 h p.i. (Fig. 1G).

In order to investigate the relative contribution of infected versus uninfected cells on *E_GSH_* of macrophage population, we assessed the biosensor response of bystander and *Mtb* infected U937 macrophages. *Mtb* cells stained with a lipophilic dye PKH26 were used to infect U937/Grx1-roGFP2 at moi 10. About 42.75%±4.05 of U937/Grx1-roGFP2 cells were infected with PKH26-labeled *Mtb* (PKH^+ve^/GFP^+ve^) (Fig. 2A). Interestingly, bystander U937 (PKH^−ve^/GFP^+ve^) showed a greater 405/488 ratio as compared to infected cells (PKH^+ve^/GFP^+ve^) at each time point tested (Fig. 2B). At 24 h p.i., *E_GSH_* of bystander cells was −276 ± 2 mV as compared to *E_GSH_*of −286 ± 2 mV in the case of *Mtb* infected cells (Fig. 2B). This suggests that the redox physiology of infected and bystander macrophages are distinctly affected during *Mtb* infection. Additionally, U937/Grx1-roGFP2 cells infected with *Mtb* genetically expressing red-fluorescent protein (RFP:tdTomato) confirmed a higher 405/488 ratio of bystander cells (RFP^−ve^/GFP^+ve^) as compared to infected cells (RFP^+ve^/ GFP^+^) (Fig. 2C). Infection of U937/Grx1-roGFP2 with PKH labeled heat-killed *Mtb* (*Hk*-*Mtb*) did not increase oxidative stress in the infected or bystander cells (Fig. 2D), indicating that processes such as secretion of bioactive lipids or proteins are the likely modulators of intramacrophage *E_GSH_*. The secretory proteins of Esx-1 family of *Mtb* are known to induce oxidative stress in the infected macrophages [25]. Agreeing to this, the *Mycobacterium bovis* (BCG) strain, defective in secreting Esx-1 proteins, elicits marginal degree of biosensor oxidation in both infected or bystander U937/Grx1-roGFP2 (Fig. 2E). Viable *Mtb*, *Hk*-*Mtb*, and BCG strains were internalized to a comparable degree by U937 (Fig. S1), precluding the influence of variations in the initial uptake rates on the *E_GSH_* of macrophages.

**Fig. 2.**
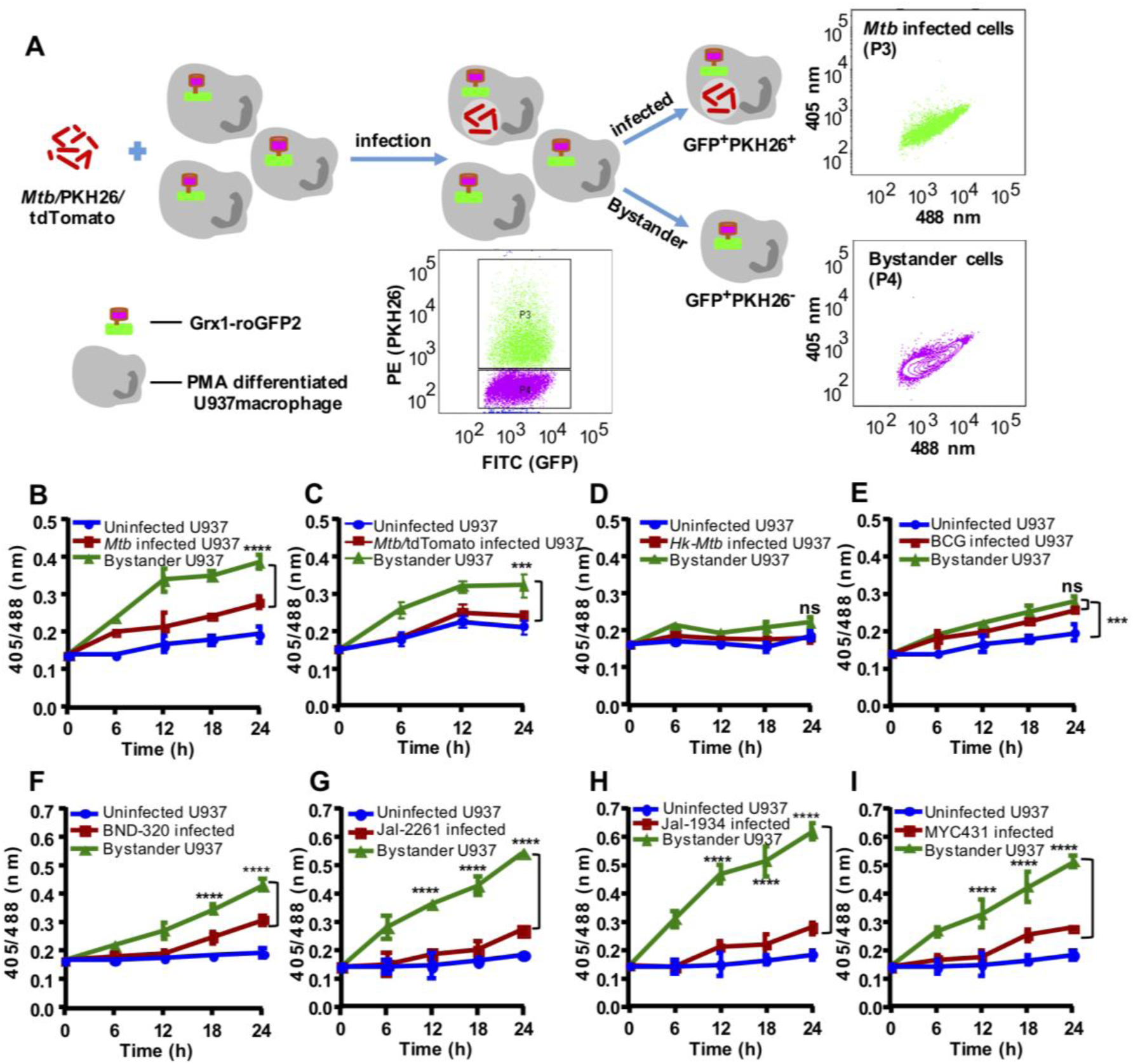
*Mycobacterium tuberculosis (Mtb)* induces higher oxidative shift in *E_GSH_* of bystander as compared to infected U937 macrophages (Mφ). **(A)** The *Mtb* H37Rv bacilli were stained with a membrane staining dye PKH26 and PMA differentiated U937/Grx1-roGFP2 cells were infected at MOI 10. The sensor response was measured by flow cytometry. Based on the PKH26 fluorescence emitted by *Mtb* inside Mφ, the U937/Grx1-roGFP2 Mφ were gated into infected (P3) or bystander subpopulations (P4) and the dot plot spectra of *Mtb* infected and bystander U937/Grx1-roGFP2 at 24 h is shown. **(B)** The line graph showing biosensor response at various time points in case of uninfected, *Mtb* infected, and bystander U937/Grx1-roGFP2. (C) *Mtb* stably **(B)** expressing red fluorescent protein, tdTomato (*Mtb*/tdTomato) was used to infect U937/Grx1-roGFP2 (MOI 10) and the biosensor response of uninfected, infected and bystander cells was measured over time. Biosensor response of infected and bystander U937/Grx1-roGFP2 cells upon infection with *PKH-26* labeled **(D)** heat-killed *Mtb* (Hk-*Mtb*), (E) *M. bovis* BCG strain, drug resistant clinical isolates of *Mtb* (F) BND-320, **(G)** Jal-2261, **(H)** Jal-1934 and **(I)** MYC-431. Error bars represent standard deviations from the mean. ***p<0.001; **** p<0.0001 (two-way ANOVA). Data are representative of at least three independent experiments performed in triplicate.

Since HIV-1 infected individuals are frequently co-infected with the drug-resistant strains of *Mtb* [26, 27], we assessed the impact of single drug-resistant (SDR; BND320), multi-drug resistant (MDR; Jal2261 and Jal 1934) and extensively drug-resistant (XDR; Myc 431) strains of *Mtb* isolated from patients [28] on *E_GSH_* of U937. Infection with BND320 induces an increase in 405/488 ratio of infected and bystander macrophages to a degree comparable to *Mtb* H37Rv (Fig. 2F). However, infection with MDR and XDR strains stimulated a significantly higher oxidative shift in *E_GSH_* of bystander cells as compared to infected U937 (Fig. 2G, 2H, and 2I), and also as compared to bystander cells in case of infection with *Mtb* H37Rv (compare Fig 2B with Fig 2G, 2H, and 2I). Altogether, these results confirm that infection with *Mtb* drives changes in *E_GSH_*of infected and bystander U937 cells and that clinical drug-resistant isolates are potent inducers of oxidative stress in macrophages.

### Reactivation of HIV-1 upon co-culturing with *Mtb* Infected macrophages

HIV-1 infects multiple immune cells including macrophages, lymphocytes, and dendritic cells, whereas macrophages are the major host cells for *Mtb*. Therefore, how *Mtb* infected macrophages communicates with HIV-1 infected cells of different origin remain unknown. The induction of oxidative stress in bystander cells raised a possibility that *Mtb* infected macrophages could reactivate virus by modulating redox physiology of neighboring cells chronically infected with HIV-1. This communication can be mediated either by direct cell-cell contact or often by bioactive soluble factors (*e.g.,* cytokines). To assess these possibilities, we set up a co-culturing experiment wherein *Mtb* infected U937 macrophages were cultured with J-Lat 10.6 lymphocytes latently infected with HIV-1. The latent HIV-1 can be reactivated by various stimuli such as phorbol esters [12-Otetradecanoylphorbol-13-acetate (TPA)], prostratin, and TNFα [29]. The integrated HIV-1 genome encodes GFP, which allows precise quantification of HIV-1 reactivation using flow cytometry. As expected, pretreatment of J-Lat with TNFα (10 ng/ ml) induced significant HIV-1 reactivation, which translated into a greater percentage of GFP+ cells over time (Fig. S2A). Co-culturing of *Mtb* infected U937 with J-Lat also showed a time-dependent increase in GFP+ cells, indicating HIV-1 reactivation (Fig. S2A). However, co-culturing with either uninfected or *Hk-Mtb* infected U937 significantly attenuated the induction of GFP from J-Lat cells (Fig. S3A). As an additional verification, we examined the effect of co-culturing on a monocytic cell line (U1) of HIV-1 latency [30]. The U1 shows basal expression of two integrated copies of HIV-1 genome, but gene expression and viral replication can be reactivated by various stimuli such as PMA, TNFα, IFN-γ, GM-CSF [31]. We tracked HIV-1 reactivation by immuno-staining for HIV-1 core protein, p24 and quantified using flow cytometer. First, we confirmed that PMA treatment reactivated HIV-1 from U1 in a time-dependent manner (Fig. S3B). Second, similar to J-Lat findings, HIV-1 reactivation was readily observed upon co-culturing of U1 cells with U937 infected with *Mtb*, whereas co-culturing with uninfected U937 macrophages or *Hk*-*Mtb* infected U937 macrophages partially reactivated HIV-1 (Fig. S2B).

To examine if this may be related to the release of soluble factors from *Mtb* infected macrophages, we treated J-LAT cells with culture supernatants derived from *Mtb* infected U937 macrophages at 24 h p.i. A time-dependent increase in GFP expression was induced by the addition of supernatant (50%, v/v) from *Mtb* infected U937 on to J-Lat cultures (Fig. 3A). Culture supernatant (50%, v/v) from uninfected U937 or *Hk-Mtb* infected macrophages showed diminished expression of GFP from J-Lat (Fig. 3A). Similar to J-Lat, treatment of U1 cells with the supernatant derived from *Mtb* infected U937 macrophages reactivated HIV-1 to the highest level as compared to supernatant from *Hk-Mtb* infected or uninfected U937 (Fig. 3B). Marginal reactivation of HIV-1 by the supernatant of uninfected U937 macrophage is perhaps due to the known effect of PMA (used as a differentiating agent) on proinflammatory cytokines secretion in the extracellular milieu [32]. Agreeing to this, supernatant from undifferentiated U937 monocytes completely failed to resuscitate HIV-1 from U1 and J-LAT (Fig. 3A and 3B). To decisively rule out the influence of PMA on HIV-1 reactivation in our assays, we infected RAW264.7 murine macrophages with *Mtb* and collected supernatant at 24 h p.i. As a control, the supernatant was also collected from the uninfected RAW264.7 macrophages. Addition of the supernatant from uninfected RAW264.7 was completely ineffective in HIV-1 reactivation from U1 cells, whereas supernatant derived from *Mtb* infected RAW264.7 was fully potent in reactivating HIV-1 (Fig. 3C).

**Fig. 3.**
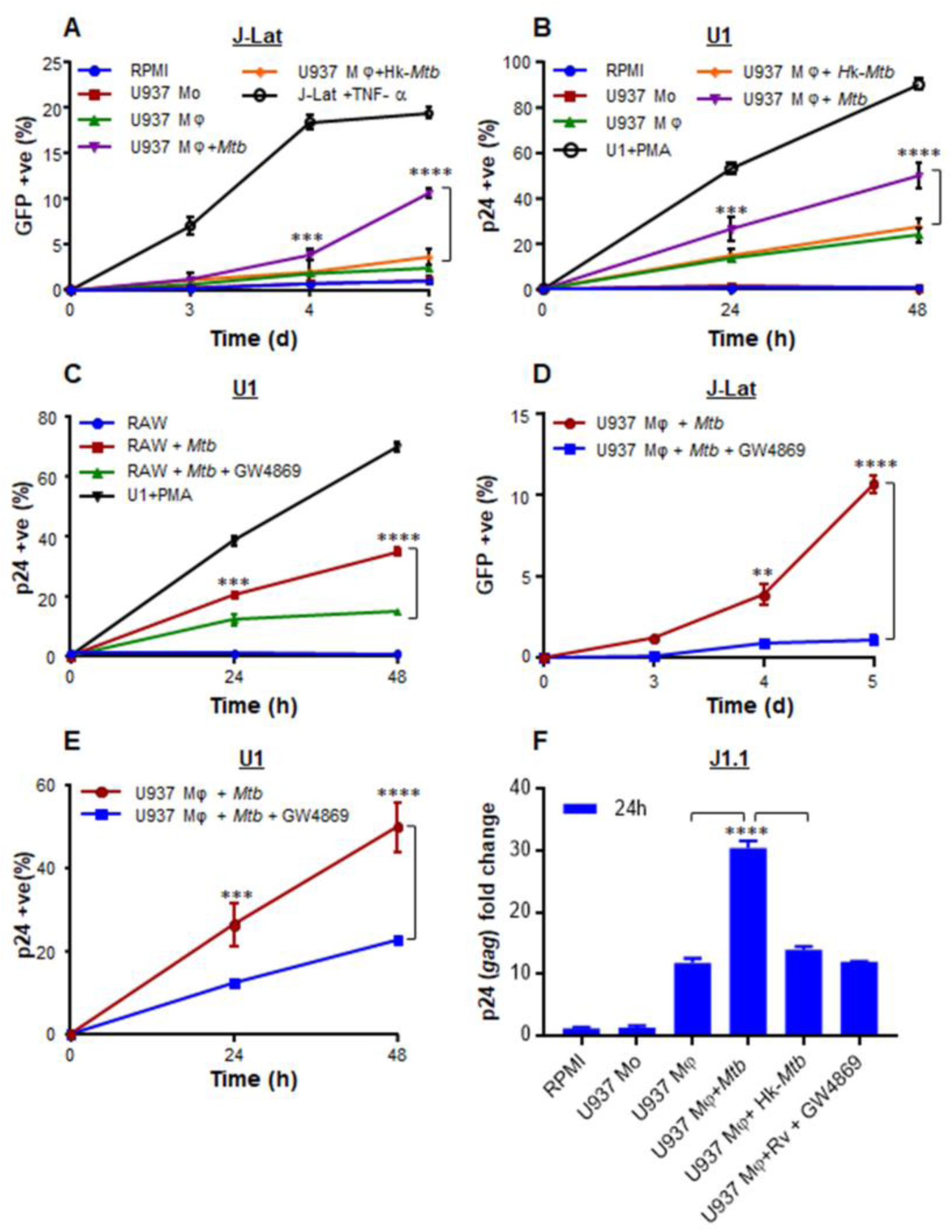
Culture supernatant derived from *Mtb* infected macrophages (Mφ) reactivates HIV-1. We determined the influence of culture supernatant derived from *Mtb* infected Mφ on three latent cell lines of HIV-1: J-Lat, U1, and J1.1. The U937 Mφ were infected with *Mtb* or Hk-*Mtb* (MOI 10). At 24 h p.i., culture supernatant from the infected macrophages was collected, passed through 0.2 µm filter, and diluted in fresh RPMI medium (1:1, v/v). This supernatant was used to culture J-Lat and U1 cells and HIV-1 reactivation was monitored over time by measuring **(A)** GFP fluorescence in J-Lat and by **(B)** p24 immuno-staining in U1 cells. As controls, we similarly monitored HIV-1 reactivation from J-Lat and U1 cells, cultured in the supernatant derived from U937 monocytes (mo) and PMA-differentiated U937 Mφ. HIV-1 reactivation upon treatment of J-Lat and U1 with TNFα (10 ng/ml) and PMA (5 ng/ml), respectively, was taken as positive control. **(C)** HIV-1 reactivation in U1 upon treatment with the supernatant derived from RAW264.7 Mφ infected with *Mtb* for 24 h. Supernatant from uninfected RAW264.7 Mφ and PMA-treatment were used as negative and positive controls for HIV-1 reactivation, respectively. To investigate if supernatant mediated HIV-1 reactivation is dependent on the presence of extracellular vesicles (*e.g.*, exosomes), we treated *Mtb* infected U937 and RAW264.7 Mφ with an inhibitor of exosome secretion (GW4869) for 24 h, followed by supernatant collection and reactivation of HIV-1 in U1 **(C and E)**, J-Lat **(D)**, and J1.1 **(F)** cells as described earlier. HIV-1 reactivation was measured by p24 (*gag*) qRT-PCR in J1.1 cells. Error bars represent standard deviations from the mean. ** p<0.01; *** p<0.001; ****p<0.0001. Data are representative of at least two independent experiments performed in triplicate.

### Exosomes derived from *Mtb* infected macrophages and mice reactivate HIV-1

Exosomes released from *Mtb* infected macrophages have been shown to direct NFκB-mediated production of proinflammatory cytokines and chemokines from bystander cells [33]. Since HIV-1 reactivation is fundamentally dependent on NFκB and proinflammatory cytokines/chemokines [34], we tested the role of exosomes in HIV-1 reactivation. To begin understanding this, we infected U937 and RAW264.7 macrophages with *Mtb* and treated infected macrophages with a well-known inhibitor of exosome secretion, GW4869 [35]. At 24 h post-treatment with GW4869, we collected supernatant and performed HIV-1 reactivation in J-Lat and U1 as described earlier. As shown in figure 3D and 3E, HIV-1 reactivation in J-Lat and U1 was observed in case of supernatant derived from *Mtb* infected macrophages, whereas a significant reduction in HIV-1 reactivation was detected in case of supernatant derived from GW4869 treated *Mtb* infected macrophages (Fig. 3D and 3E). Furthermore, we have taken a third cell line (J1.1 lymphocyte) latently infected with HIV-1 [36] and confirmed that only the supernatant derived from *Mtb* infected macrophages reactivated HIV-1 (as shown by *gag* qRT-PCR) as compared to supernatant from uninfected, *Hk-Mtb*, and GW4869 treated macrophages (Fig. 3F). These results suggest a role for exosomes secreted by *Mtb* infected macrophages in reactivating HIV-1. Since supernatant-derived from *Mtb* infected U937 and RAW264.7 macrophages resulted in an identical degree of HIV-1 reactivation, further experiments are conducted using RAW264.7 cells only. This was necessary to rule out any artifactual influence of PMA used to differentiate U937 monocytes prior to infection with *Mtb*. Furthermore, exosomes derived from *Mtb* infected RAW264.7 have been shown to possess immune-modulatory properties comparable to exosomes isolated from the serum of *Mtb* infected mice or humans [37].

We infected RAW264.7 with *Mtb* H37Rv, and at 24 h p.i., exosomes were isolated. Transmission electron microscopy (TEM) confirmed the size of exosomes to be 80-150 nm [n= 100] (Fig. 4A), which is consistent with the previous studies [38]. Immuno-gold labeling with CD63 primary antibody followed by TEM confirmed the presence of this classical exosomal marker on the surface (Fig. 4B) [39]. Immuno-blot analysis for exosome-specific markers such as Rab5b, Alix, and CD63 established their enrichment on the exosomal fraction relative to cell lysate (Fig. 4C). LAMP2 was present in both the fractions as previously shown (Fig. 4C) [39]. Finally, we demonstrated that the pretreatment of *Mtb* infected RAW264.7 macrophages with GW4869 significantly reduced the release of exosome as revealed by the loss of CD63 marker as compared to untreated or *Mtb* infected RAW264.7 macrophages (Fig. 4D). Altogether, we confirmed the isolation of exosomes from *Mtb* infected macrophages for investigating redox-dependent activation of HIV-1.

**Fig. 4.**
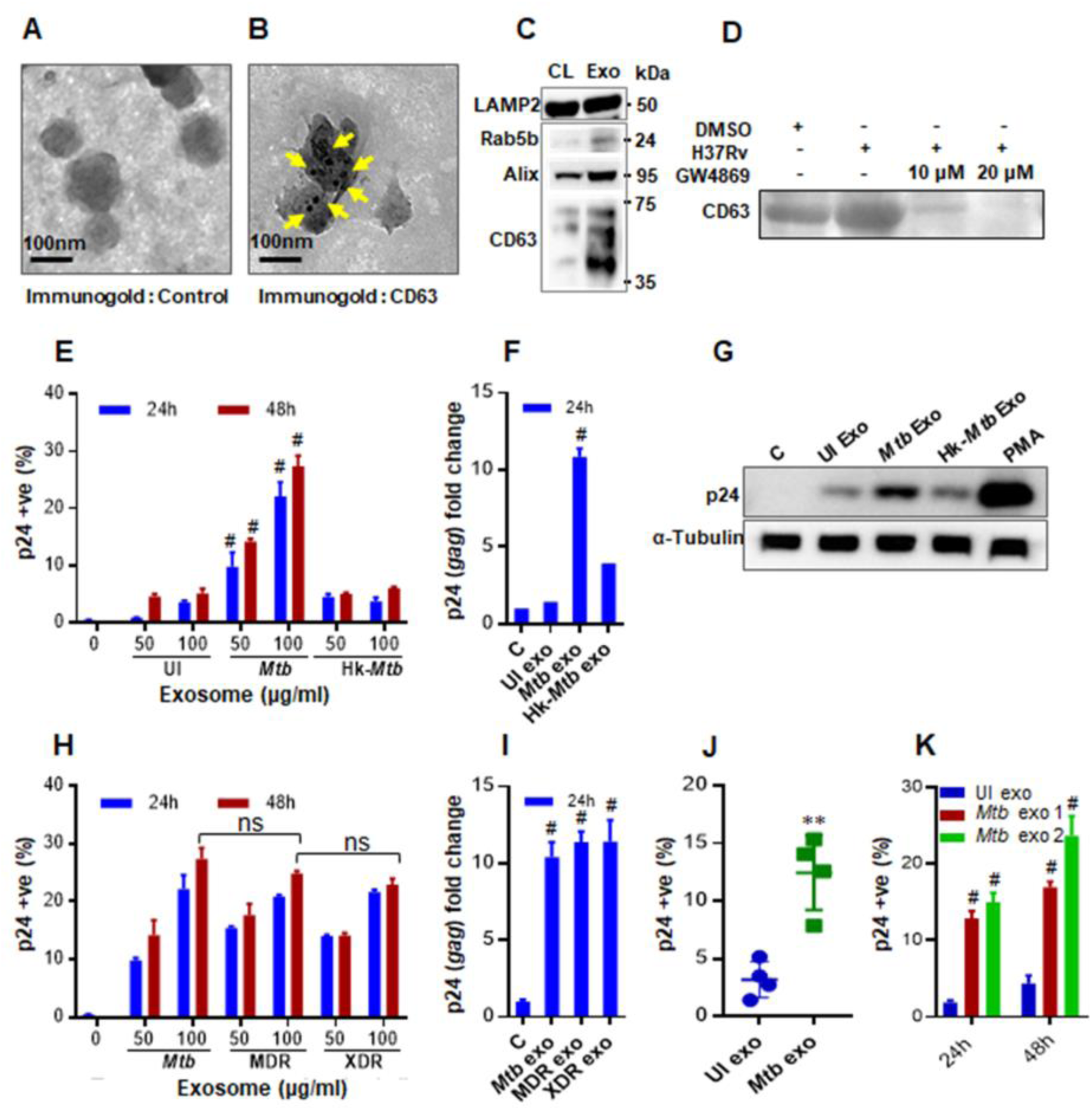
Exosomes derived from *Mtb* infected macrophages (Mφ) and mice induce HIV reactivation. Exosomes were isolated from enriched cell culture supernatant of *Mtb* infected RAW 264.7 Mφ. Exosomes were purified by using the ExoQuick precipitation solution and morphologically characterized by transmission electron microscopy (TEM). **(A)** Control, **(B)** Immunogold labeling of exosomes using the antibody against exosomes surface marker, CD63. Scale bar, 100 nm, **(C)** Immuno-blot analysis of LAMP2, Rab5b, Alix and CD63 in cell extract (CL; 30 μg) and purified exosomes (Exo, 30 μg) from *Mtb* infected RAW264.7 Mφ, **(D)** Immuno-blot analysis of exosome-specific marker CD63 to show dose dependent reduction in exosome production by *Mtb* infected RAW 264.7 Mφ upon treatment with GW4869. Exosomes were derived from equal number of RAW 264.7 Mφ in each case and equivalent volume was loaded in each lane. **(E)** U1 cells were treated with 50 and 100 μg/ml of purified exosomes and HIV-1 reactivation was measured by flow cytometry using PE labeled antibody specific to p24 (Gag) antigen at the indicated time-points, **(F)** qRT-PCR for *gag* transcript at 24 h post-treatment with 100 μg/ml of purified exosomes, **(G)** immuno-blot for p24 in the cell lysate (50 μg) at 24 h post-treatment with 100 μg/ml of purified exosomes. CTR and UI exo denote HIV-1 reactivation without any treatment or upon treatment with exosomes (100 μg/ml) derived from uninfected RAW 264.7 Mφ, respectively. RAW264.7 Mφ were infected with MDR (Jal-1934) and XDR (Myc431) strains for 24 h and exosomes were isolated. U1 cells were treated with exosomes and HIV-1 reactivation was monitored at the indicated time by flow cytometry using **(H)** PE labeled antibody specific to p24 (Gag) antigen and **(I)** qRT-PCR of *gag* transcript. BALB/c mice were infected with 100 CFU of *Mtb* H37Rv or were left uninfected. Exosomes were isolated from mouse serum and from the lung tissue at 20 weeks post infection using ExoQuick and quantified by micro BCA assay. U1 cells were treated with exosomes derived from **(J)** serum (2 mg/ml) or **(K)** from lungs (*Mtb* exo1 - 100 μg/ml, *Mtb* exo2 - 200 μg/ml) and HIV reactivation was measured by flow cytometry using fluorescent tagged antibody (PE labeled) specific to p24 (Gag) antigen. Error bars represent standard deviations from the mean. Statistical analyses were performed using two tailed unpaired t-test **(J)**, one-way ANOVA **(F, I)** and two-way ANOVA **(E, H, K)**. ns-non significant; **p<0.01; # p<0.001. Data are representative of at least two independent experiments performed in duplicate.

Various concentrations of exosomes isolated from *Mtb* infected RAW264.7 macrophages were used to treat U1 and HIV-1 reactivation was measured by p24 immuno-staining and qRT-PCR of *gag* transcript. As compared to uninfected control, *Mtb* specific exosomes significantly induce HIV-1 reactivation from U1. Both p24 staining and *gag* qRT-PCR displayed ∼ 4-fold increase as compared to uninfected control (Fig. 4E, 4F, and 4G). Exosomes isolated from macrophages infected with *Hk*-*Mtb* were ineffective in reactivating HIV-1 (Fig. 4E, 4F, and 4G). However, the exosomes derived from RAW264.7 macrophages infected with MDR and XDR strains of *Mtb* were equally potent in reactivating HIV-1 (Fig. 4H and 4I). Finally, to conclusively show that exosomes derived from *Mtb* infection reactivate HIV-1, we treated U1 with exosomes isolated from serum and lungs of mice chronically infected with *Mtb* and stained for p24. In both cases, we observed a significant reactivation (∼ 5-fold) of HIV-1 as compared to exosomes isolated from uninfected animals. (Fig. 4J and 4 K) However, exosomes isolated from *Mtb* infected lungs reactivated HIV-1 at a lower concentration than serum (Fig. 4J and 4 K). Taken together, our data suggest that exosomes could be one of the important mediators of HIV-1 reactivation during *Mtb* infection.

### Exosomes from *Mtb* infected macrophages modulate redox potential and host gene expression of U1 cells

Elevated ROS, RNI, and depletion/oxidation of intracellular thiols (cysteine, thioredoxin, GSH) were shown to activate the HIV-1 long terminal repeat (LTR) through the redox-responsive transcription factor NFκB [40, 41]. Therefore, we first tested if exosomes derived from RAW 264.7 infected with various *Mtb* strains induce oxidative stress to reactivate HIV-1. Treatment of U1/Grx1-roGFP2 with exosomes derived from RAW 264.7 infected with *Mtb* H37Rv, MDR, and XDR strains uniformly increases biosensor ratio at 24 h and 48 h post-treatment, indicating oxidative stress (Fig. 5A). As expected, exosomes derived from uninfected or *Hk*-*Mtb* infected RAW 264.7 failed to induce oxidative stress in U1 cells (Fig. 5A). To show that exosomes-triggered oxidative shift in *E_GSH_*precedes HIV-1 reactivation, we pretreated U1 cells with the GSH-specific antioxidant, N-acetyl cysteine (NAC), followed by exosomes addition. Treatment with NAC entirely abrogated the potential of *Mtb* specific exosomes to reactivate HIV-1 in a concentration dependent manner (Fig. 5B). This result reiterates that oxidative stress is likely to be an important mechanism induced by *Mtb*-specific exosomes to reactivate HIV-1.

**Fig. 5.**
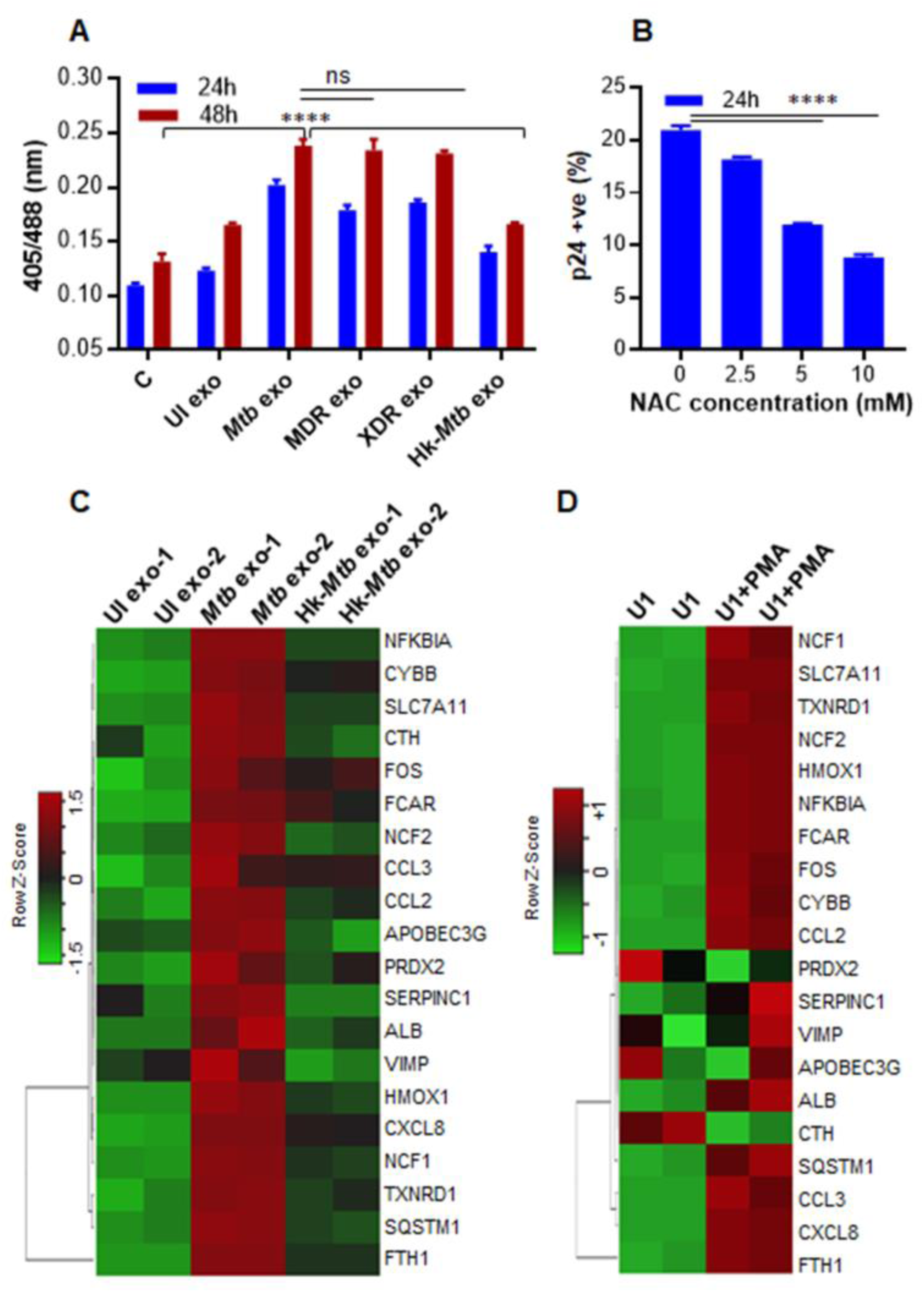
*Mtb* specific exosomes reactivate HIV-1 by inducing oxidative stress. **(A)** RAW264.7 Mφ were infected with *Mtb*, Hk-*Mtb*, Jal-1934 (MDR) and MYC431 (XDR) at moi 10. At 24 h p.i., exosomes were isolated from the culture supernatant, and 100 μg/ml of exosomes were used to treat U1/Grx1-roGFP2 for 24 and 48 h. Ratiometric biosensor response was measured by flow cytometry. UI exo: denotes exosomes isolated from uninfected RAW264.7. **(B)** U1 cells were pre-treated with N-acetyl cysteine (NAC) for 1 h at indicated concentrations followed by exposure to *Mtb* specific exosomes (100 μg/ml) for 24 h. HIV reactivation was measured by flow cytometry using fluorescent tagged antibody (PE labeled) specific to p24 (Gag) antigen. Data are representative of at least three independent experiments performed in triplicate. **(C)** U1 cells were treated with exosomes (100 μg/ml) isolated from uninfected*, Mtb* infected, and *Hk-Mtb* infected RAW264.7. Total RNA from U1 cells was isolated after 12 h of exosome treatment, followed by expression analysis of 185 genes specific to HIV host response and oxidative stress response utilizing NanoString customized gene expression panels. Data were normalized by nSolver (NonoString) software. Genes with significant changes were selected based on p<0.05 and fold change >1.5 between *Mtb versus* UI, *Mtb versus* Hk-*Mtb*, Hk-*Mtb versus* UI. Data is displayed as heat maps of three groups together. **(D)** Expression profile of HIV host response and oxidative stress response in U1 monocytes and U1 macrophages (PMA-differentiated). Error bars represent standard deviations from the mean. ****p<0.0001 (two-way ANOVA). Data are representative of at least two independent experiments performed in duplicate.

To further understand the mechanism of exosomes mediated HIV-1 reactivation, we examined the influence of *Mtb* specific exosomes on host gene expression using NanoString nCounter gene expression analysis. This technology permits for the absolute quantification of a large number of RNA transcripts without any requirements for reverse transcription or DNA amplification [42]. From the NanoString panel, we focused on host genes that respond to HIV replication (*e.g.,* HIV receptors-ligands, proteins involved in HIV replication, inflammatory response, apoptosis, cell cycle, and transcription activators of HIV LTR) as well as genes involved in oxidative stress response (Table S1a-S1c). Exosomes derived from uninfected or *Mtb* (live or *Hk*) infected RAW 264.7 macrophages were used to treat U1 and total RNA was isolated at 12 h post-treatment. An early time point is taken to ensure that we capture gene expression changes that precede HIV-1 reactivation upon exosomes challenge. Also, at a later time point, the primary effect of exosomes can be masked due to transcriptional changes in response to HIV proliferation and associated cytopathic consequences. PMA treated U1 were taken as a positive control. Consistent with our biosensor data, treatment with viable *Mtb*-specific exosomes induces genes encoding components of superoxide-producing enzyme-NADPH oxidase (*e.g.,* cytochrome B-245 Beta chain [CYBB], NCF1, and NCF2) (Fig. 5C). As a consequence, several genes involved in mitigating ROS and maintaining redox balance such as PRDX2 (peroxiredoxin) [43], TXNRD1 (thioredoxin reductase 1) [44], sodium-dependent cysteine-glutamate antiporter (SLC7A11) [45], and ferritin (FTH1) [46] were highly induced in U1 treated with *Mtb*-exosomes than *Hk*-*Mtb* or uninfected exosomes (Fig. 5C). Additionally, a gene encoding heme oxygenase 1 (HMOX1) is highly induced in U1 treated with *Mtb*-exosomes (Fig. 5C). HMOX1 is involved in heme catabolism and its dysregulation and polymorphism in its promoter are linked with HIV-associated neurocognitive disorder [47]. Gene encoding a member of the selenoprotein family (VCP-interacting membrane protein [VIMP], also known as selenoprotein S was induced by *Mtb*-exosomes (Fig. 5C). VIMP is a redox-sensing protein that regulates inflammation by mediating cytokine production [48], suggesting that VIMP can coordinate HIV-1 reactivation by triggering an inflammatory reaction in response to oxidative stress induced by *Mtb*-exosomes.

In agreement with HIV-1 reactivating potential of *Mtb*-exosomes, genes involved in ensuring HIV-1 LTR activation were induced. For example, NFκBIA and SQSTM1/P62 involved in the activation of transcription factor NFκB were upregulated [49](Fig. 5C). Another transcription factor FOS, which reactivates HIV-1 [50], was also upregulated by *Mtb*-exosomes. Several genes encoding host inflammatory mediators were induced by *Mtb*-exosomes. For examples, expression of CXC chemokine subfamily (CXCL8), monocyte chemoattractant protein 1 (MCP1; CCL2) and macrophage-inflammatory-protein-1 alpha (MIP-1a; CCL3) showed greater expression in U1 treated with *Mtb* specific exosomes as compared to *Hk-Mtb* or uninfected exosomes (Fig. 5C). All of these factors are well known to facilitate HIV-1 infection and promote replication in macrophages [51]. In facts, higher levels of CCL2 were detected in the bronchoalveolar lavage (BAL) fluid of pulmonary TB patients and pleural fluid of HIV-1 infected patients, indicating the importance of this proinflammatory cytokine in HIV-TB co-infection [52]. Interestingly, apolipoprotein B mRNA-editing enzyme catalytic polypeptide-like 3G (APOBEC3G), a known HIV restriction factor [53], was also induced by *Mtb*-exosomes. In addition to viral restriction, APOBEC3G activity also promotes heterogeneity in HIV sequence resulting in HIV phenotypes with greater capacity to escape immune pressures [54]. Therefore, it’s likely that co-infection with *Mtb* and subsequent exosomes release can accelerate APOBEC3G mediated generation of more fit viral variants. Consistent with this, pleural fluid mononuclear cells (PFMCs) from HIV/TB coinfected patients show higher expression of APOBEC3G [55]. Lastly, gene encoding serpin peptidase inhibitor, clade C (SERPINC1; antithrombin) that induces at very early stages of HIV-1 replication, and is a component of the viral core [56], was upregulated by *Mtb* exosomes (Fig. 5C). We found a striking similarity between the transcriptional signatures of host genes induced by *Mtb*-exosomes to those induced by the PMA treatment (Fig. 5D), confirming that *Mtb*-exosomes are eliciting host response associated with HIV-1 reactivation. Taken together, exosomes released by *Mtb* infected macrophages modulate host redox metabolism and inflammatory response to ensure HIV-1 reactivation.

### Influence of *Mtb*-specific exosomes on oxidative phosphorylation (OXPHOS) of U1 cells

We have previously shown that HIV-1 reactivation precedes an oxidative shift in mitochondrial *E_GSH_* and the mitochondrial redox physiology closely coincides with the progression of HIV disease [20]. This indicates that *Mtb*-exosomes might influence the mitochondrial physiology of HIV infected cells to promote oxidative stress and viral replication. To examine this possibility, we performed a real-time assessment of the mitochondrial function of U1 cells in response to *Mtb*-specific exosomes. We quantified mitochondrial physiology by measuring several key parameters associated with oxidative phosphorylation (OXPHOS) using Seahorse XF Flux technology (Fig. 6A).

**Fig. 6.**
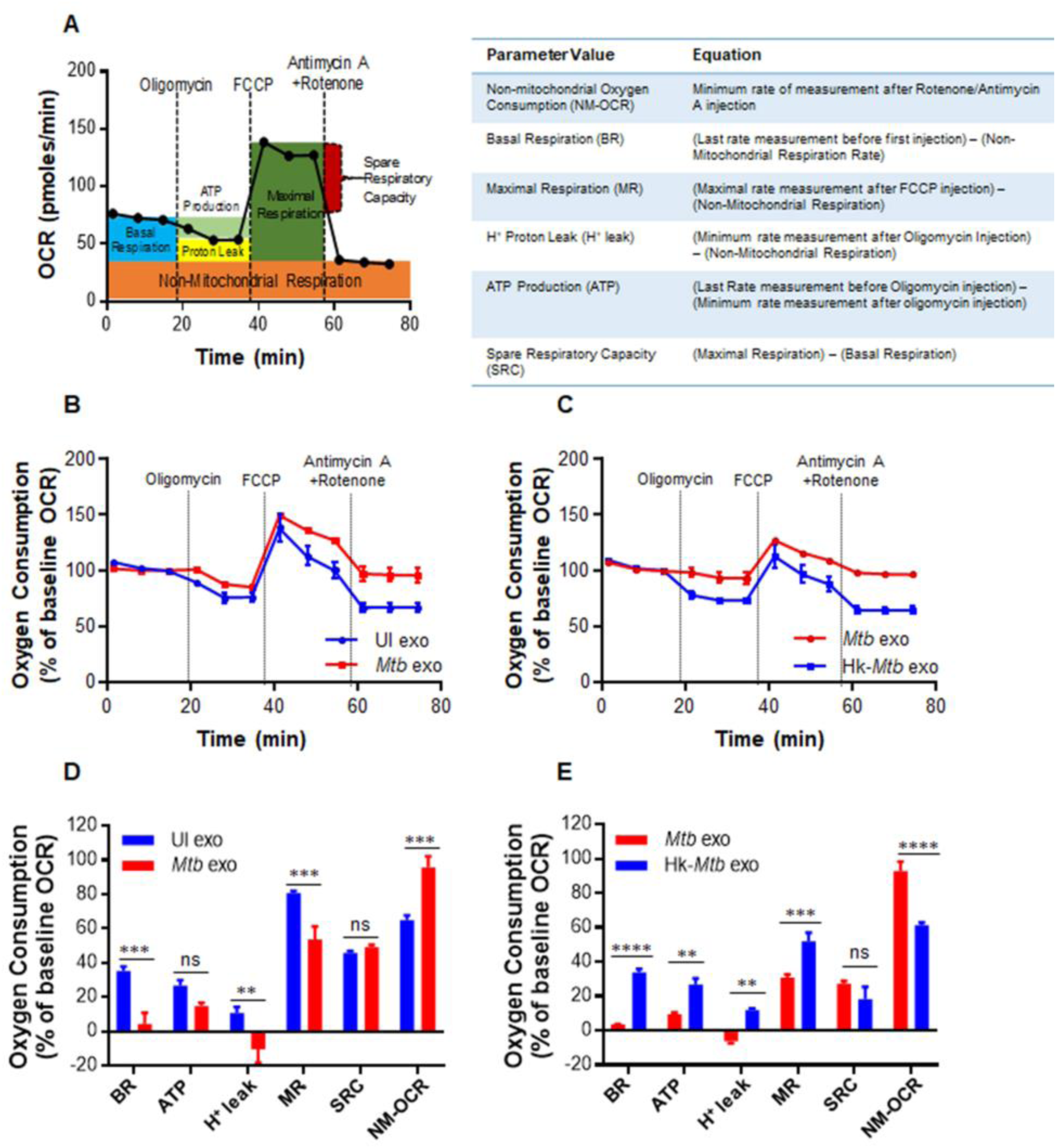
*Mtb*-specific exosomes modulate OXPHOS of U1 cells. **(A)** Schematic presentation of Agilent Seahorse XF Cell Mito Stress test profile of the key parameters of mitochondrial respiration. **(B and C)** RAW264.7 macrophages were infected with *Mtb* and Hk-*Mtb* at moi 10. At 24 h p.i., exosomes were isolated from the culture supernatant, and 100 μg/ml of exosomes were used to treat U1 for 48 h. Respiratory profile of U1 cells on treatment with exosomes isolated under indicated conditions. UI exo: exosomes isolated from uninfected RAW264.7. Respiratory profile was measured and expressed as percent of basal oxygen consumption rate (%OCR). Oxygen consumption was measured without any inhibitor (basal respiration), followed by OCR change upon sequential addition of oligomycin (1 μM; ATP synthase inhibitor) and cyanide-4 [trifluoromethoxy]phenylhydrazone (FCCP; 0.25 μM), which uncouples mitochondrial respiration and maximizes OCR. Finally, respiration was completely shut down by inhibiting respiration using antimycin A and Rotenone (0.5 μM each), inhibit complex III and I, respectively. **(D and E)** Various respiratory parameters as outlined in the table of panel A were measured using the values obtained from the dataset depicted in panel **(B)** and **(C)** and as described in materials and methods. Error bars represent standard deviations from the mean. ns-non significant; **p<0.01; ***p<0.001 (two tailed unpaired t-test). Data are representative of at least two independent experiments performed in triplicate.

Several parameters of mitochondrial respiration, including basal respiration (Basal-Resp), ATP-linked respiration, proton leak and spare respiratory capacity (SRC), were derived by the successive addition of pharmacological agents to the exosomes challenged U1 cells, as outlined in Figure 6A. To determine each parameter, three reiterated rates of oxygen consumption (OCR) are made over an 80-minute period. Firstly, baseline cellular oxygen consumption (OCR) is measured, from which Basal-Resp is derived by subtracting non-mitochondrial respiration. Secondly, an inhibitor of complex V (oligomycin) is added, and the resulting OCR is used to calculate ATP-linked OCR (by deducting the OCR after oligomycin addition from baseline cellular OCR) and proton leak (by subtracting non-mitochondrial respiration from the OCR upon oligomycin addition). Thirdly, maximal respiration (Max Resp), which is the change in OCR after uncoupling ATP synthesis from electron transport by adding carbonyl cyanide-4-(trifluoromethoxy)phenylhydrazone (FCCP). Lastly, antimycin A, a complex III inhibitor, and rotenone, a complex I inhibitor, are added together to inhibit ETC function, revealing the non-mitochondrial respiration (Non-Mito Resp). The spare respiratory capacity (SRC) is calculated by subtracting basal respiration from maximal respiratory capacity.

U1 cells were treated with exosomes isolated from RAW264.7 infected with viable or *Hk*-*Mtb* for 48 h. Following this, U1 cells were seeded on to the XF cartridge plates and subjected to the mitochondrial stress test to measure various OXPHOS parameters as described earlier. As compared to exosomes from uninfected macrophages, treatment of U1 cells with *Mtb*-specific exosomes significantly decreased various respiratory parameters including basal respiration, ATP-linked OCR, and H^+^ leak (Fig. 6B and 6D). In contrast, the non-Mitochondrial Resp was significantly increased, whereas SRC was not significantly affected (Fig. 6B and 6D). Exosomes derived from macrophages infected with Hk-*Mtb* modulate OXPHOS parameters comparable to exosomes from uninfected macrophages (Fig. 6C and 6E). The contrasting influence of viable and *Hk*-*Mtb* exosomes on Basal-Resp, ATP-linked OCR and proton leak indicate a profound deceleration of respiration of U1 upon reactivation of HIV-1 by *Mtb* exosomes. Generation of ROS (e.g., superoxide) is an inevitable consequence of normal mitochondrial respiration. Since *Mtb* exosomes induce an oxidative shift in *E_GSH_* of U1, a decrease in Basal-Resp and ATP-linked OCR could be a cellular strategy to avoid overwhelming oxidative stress. This would ensure successful HIV-1 reactivation without triggering the detrimental effects of ROS on U1. Agreeing to our findings, active HIV-1 replication depends largely on increased glycolytic flux rather than OXPHOS to meet the surge in biosynthetic and bioenergetics demand [57].

Our OXPHOS results also provide clues about the mechanism of an oxidative shift in *E_GSH_* upon HIV reactivation by exosomes. We observed a significant increase in non-mitochondrial O_2_ consumption in the case of U1 treated with *Mtb* specific exosomes (Fig. 6D and 6E). Non-mitochondrial O_2_ consumption is usually due to activities of enzymes associated with inflammation such as lipoxygenase, cyclo-oxygenase, and NADPH oxidase [58]. Consistent with this, our NanoString data showed an increased expression of genes encoding various components of the NADPH oxidase complex. Altogether, HIV-1 reactivation by *Mtb*-specific exosomes was associated with a marked change in OXPHOS parameters including a reduced basal OCR and ATP-linked OCR. An increased Non-Mito OCR is likely responsible for the generation of oxidative stress during HIV-TB co-infection.

### Proteomics of exosomes released upon *Mtb* infection

Having established that *Mtb* specific exosomes reactivate HIV-1 by modulating redox and bioenergetics, we sought to determine the content of exosomes. Mycobacterial components (*e.g.,* lipids, proteins, and RNA) that exert pro-inflammatory responses are consistently enriched in the exosomes of *Mtb* infected macrophages [59, 60]. However, the identity of host proteins within exosomes isolated from *Mtb* infected macrophages remains uncharacterized. This is important as immune-activated macrophages secrete several redox-signaling proteins involved in eliciting proinflammatory response and oxidative stress in the neighboring cells [43]. On this basis, we reasoned that profiling of exosome-associated host-proteins likely shed new insight on how *Mtb* induces HIV-reactivation.

We identified proteins associated with exosomes from uninfected, live *Mtb*, and *Hk Mtb* infected RAW264.7 cells using LC-MS/MS (Fig. 7A). In total, 4953 proteins were identified in all three samples with two-biological replicates (Sequest Program, Fig. 7A, Table S2). As expected, a high correlation value (> 0.90, Pearson correlation coefficient) was observed within the two biological replicates from the same group than unrelated groups (0.6-0.7) (Fig. S3C). About 80 of the 100 most identified exosomal proteins of the ExoCarta database were identified in our dataset. While ∼3250 proteins overlapped among the three groups, we discovered that 86, 298, and 142 proteins were exclusively present in the uninfected, live *Mtb,* and *Hk Mtb*, respectively (Fig. S3B). The highest numbers of proteins in case of live *Mtb* exosomes clearly indicates that *Mtb* infection promotes secretion of proteins in exosomes. Analysis of differentially expressed proteins (Table S3a-S3c) (log 2 fold, P ≤ 0.01) showed that in comparison to uninfected samples, 436 proteins were up-regulated and 290 were down-regulated in live *Mtb*-exosomes (Fig. 7B). Similarly, 390 proteins were induced and 337 were repressed in *Hk*-*Mtb* as compared to uninfected samples (Fig. 7B). Direct comparison of live versus *Hk*-*Mtb* revealed that ∼ 40 proteins were differentially enriched in exosomes derived from live *Mtb* infected macrophages (Fig. 7B). This suggests that live *Mtb*-derived exosomes contain different proteins, which might mediate more specific function such as redox imbalance and HIV reactivation upon secretion.

**Fig. 7.**
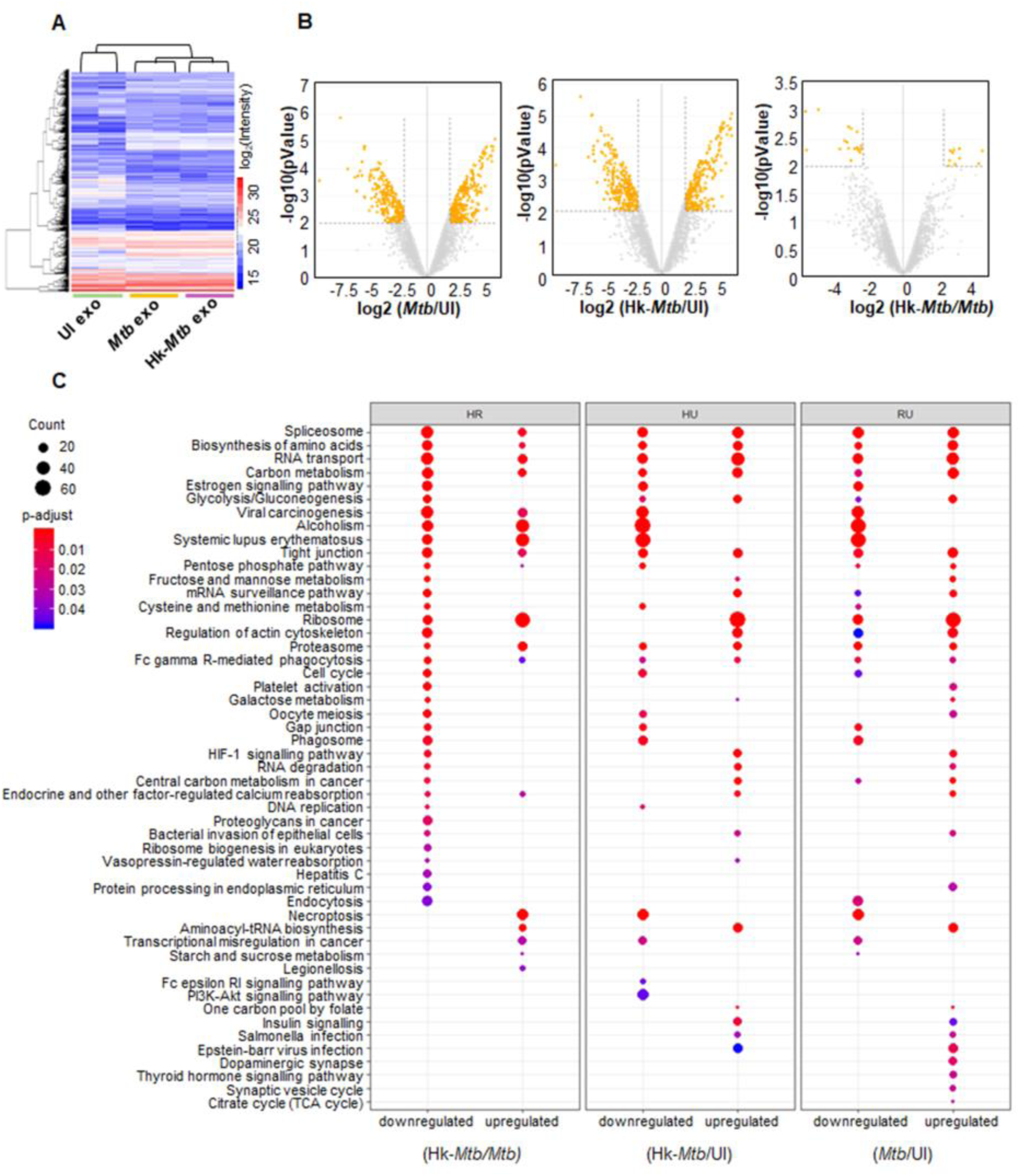
Proteomics of *Mtb* specific exosomes by LC-MS/MS. RAW264.7 Mφ were infected with *Mtb* and Hk-*Mtb* at moi 10 or left uninfected. At 24 h p.i., exosomes were isolated from the culture supernatant for LC-MS/MS. **(A)** Heat map of differentially expressed proteins of exosomes in three samples. **(B)** Volcano plots of differentially expressed proteins. Significantly up-regulated and down-regulated proteins with log2 fold change more than two are shown as orange dots. **(C)** Enriched KEGG signaling pathways, proteins upregulated or downregulated in different comparison groups: (heat killed) Hk-*Mtb* versus live *Mtb,* Hk-*Mtb* versus UI (uninfected) and live *Mtb* versus UI (uninfected).

Biological process analysis of Gene ontology (GO) showed that the differentially expressed proteins belong to diverse categories including cellular/metabolic processes, biological regulation and response to stimulus (Fig. S4). Molecular function analysis of GO revealed that most of the proteins carry out binding and catalytic activity, indicating their roles in exosomes cargo sorting, release, and uptake by the recipient cells (Fig. S5). This is further confirmed by cell component analysis wherein exosomal proteins mainly belonged to cell organelle, membrane, and macromolecular complexes involved in exosome biogenesis (Fig. S5) (Table S4a-S4c, S5a-S5c, S6a-S6c).

Next, we classified the differentially expressed proteins of each group by KEGG signaling pathway (Table S7) of KEGG database (Fig. 7C). *Mtb* exosomes were found to be enriched with proteins involved in glycolysis, gluconeogenesis, fructose and mannose metabolism, galactose metabolism, pentose phosphate pathway (PPP), cysteine and methionine metabolism. Increased oxidative stress in HIV infected patients is associated with higher glucose utilization and deficiency of cysteine and methionine [61-63]. Enrichment of sugar metabolic enzymes in *Mtb* exosomes possibly assists in HIV-1 reactivation by fueling ATP generating processes for the energy consuming functions such as virus transcription, translation, packaging, and release. Similarly, cysteine metabolism serves as a source of GSH biogenesis, while PPP enzymes provides NADPH for regenerating GSH from GSSG. Both of these activities are essential for alleviating excessive oxidative stress to avoid cell death during HIV-1 reactivation [16]. We observed enrichment of proteins coordinating RNA transport/quality control, DNA replication, and cell cycle, all of which are important for the reactivation of HIV-1 [64, 65]. Importantly, estrogen signaling proteins involved in induction of mitochondrial ROS and HIV-1 reactivation are enriched in *Mtb*-exosomes [66, 67]. HIF-1 signaling plays an important role in HIV-1 pathogenesis by facilitating viral replication and promoting lymphocyte and macrophage mediated inflammatory response [68]. HIF-1 pathway proteins are specifically induced in *Mtb* exosomes (Fig. 7C).

We discovered that several members of heat shock protein family (HSP) and galectins were exclusively enriched in *Mtb* exosomes. *Mtb* infected macrophages are reported to secrete HSPs in exosomes to induce proinflammatory response (NFκB and TNFα) from the uninfected bystander cells [69]. Furthermore, HSP such as Hsp90 potently reactivates HIV from latently infected cell lines and CD4+ T cells [70]. Likewise, galectins are consistently associated with oxidative stress, inflammation, and HIV-1 reactivation [71, 72]. Finally, STAT-1 of JAK-STAT signaling pathway, which is well known to modulate HIV replication cycles, was enriched in *Mtb*-specific exosomes [73, 74].

Altogether, infection with viable *Mtb* induces secretion of specific proteins in exosomes to reactivate HIV-1 by affecting redox, central metabolism, and inflammatory responses.

### HSPs and Galectins facilitate HIV-1 reactivation by *Mtb* exosomes

Since multiple pathways are likely to synergies during HIV-1 reactivation by *Mtb* specific exosomes, determination of the specific proteins/pathways in this process is challenging. However, to begin delineating the role of some of the components secreted in exosomes upon *Mtb* infection in HIV-1 reactivation, we decided to examine the effect of HSPs and galectins. We first examined the effect of Hsp90 pathway by using a specific inhibitor of Hsp90-17-(N-allylamino)-17-demethoxygeldanamycin (17-AAG) [70]. The inhibitor 17-AAG binds to ATPase pocket of HSP90 and compete with ATP, resulting in inactivation of chaperone function [70]. U1 cells were stimulated with PMA and HIV-1 reactivation was monitored in the presence or absence of 17-AAG by scoring for p24 positive cells using flow cytometry. The 17-AAG completely abolished PMA-mediated HIV-1 reactivation but showed no effect on the basal expression of p24 in the unstimulated cells (Fig. 8A). Similarly, 17-AAG significantly decreased HIV-1 reactivation in U1 by *Mtb*-specific exosomes (Fig. 8B).

**Fig. 8.**
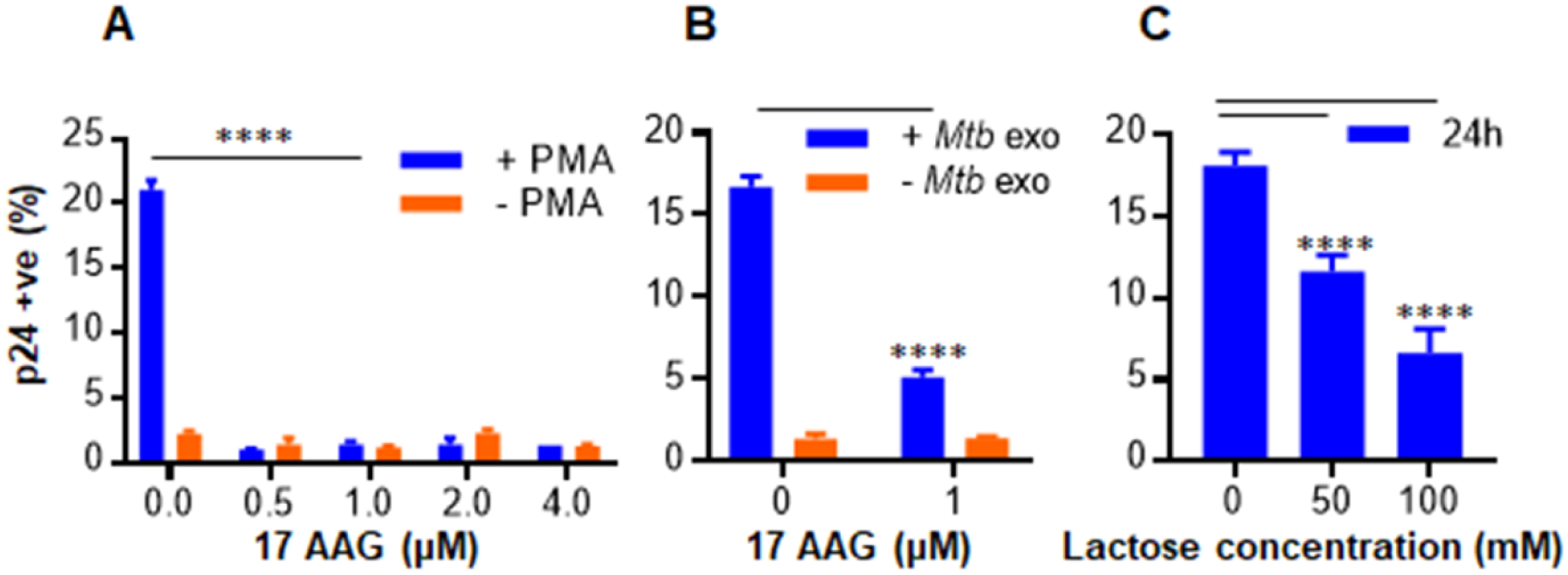
Hsp-90 and Galectin inhibitors reverse the exosomes mediated HIV-1 reactivation in U1 cells. **(A)** U1 cells were stimulated with 5 ng/ml of PMA to induce HIV reactivation in presence of indicated concentration of 17-AAG (Hsp-90 inhibitor). **(B)** U1 cells were treated with 100 μg/ml of *Mtb* exosomes isolated from RAW264.7 as described earlier and HIV reactivation was monitored in the presence of indicated concentration of 17-AAG. **(C)** *Mtb* infected RAW264.7 macrophages were treated with 50 and 100 mM of lactose (galectins inhibitor) for 24 h and exosomes were isolated. These exosomes were then used to reactivate HIV-1 from U1 cells. HIV-1 reactivation was measured by flow cytometry using fluorescent tagged antibody (PE labeled) specific to p24 (Gag) antigen. Error bars represent standard deviations from the mean. *p<0.05; **p<0.01; *** p<0.001; ****p<0.0001. One-way ANOVA **(C)** and two-way ANOVA **(A, B)**. Data are representative of at least two independent experiments performed in duplicate.

To understand how galectins modulate exosomes mediated reactivation of HIV-1, we exploited the lactose binding property of galectins. Galectins form a complex with lactose, via carbohydrate binding domain, which inhibits galectins activities [75]. Galectins control numerous biological functions mainly by binding to the cell surface associated glycoproteins or glycolipid receptors. Therefore, we thought of inhibiting the activity of galectins before it’s transported in the *Mtb*-specific exosomes to preclude its interaction with the U1 cell surface glycoproteins/receptors. To do this, we treated *Mtb* infected RAW 264.7 macrophages with 50 and 100 mM of lactose for 24 h and exosomes were isolated. These exosomes were then used to reactivate HIV-1 in U1. As shown in figure 8C, exosomes isolated from the lactose treated cells significantly reduced the ability of *Mtb*-specific exosomes to reactivate HIV-1 from U1 monocytes. Altogether, *Mtb*-specific exosomes harbor and convey cellular effectors responsible for reversing viral latency in cell lines chronically infected with HIV-1.

## Discussion

We have previously shown that a marginal oxidative shift in *E_GSH_* is sufficient to reactivate HIV-1 [20]. Others have shown that *Mtb* induces oxidative stress and GSH imbalance in the infected macrophages, animals, and humans [21, 22]. HIV-TB co-infected patients suffer from glutathione stress, metabolic deficiencies, and immune-dysfunction [21]. Critically, we now made an effort to unify in a coherent picture of these separate observations, establishing a functional link between *Mtb* induced oxidative stress and HIV-1 reactivation from latency, which may have therapeutic implications. Furthermore, we provide evidence for oxidative stress-mediated HIV-1 reactivation, which relies on the secretion of biological effectors present in the exosomes released from *Mtb* infected macrophages. Although with hindsight the connection between *Mtb*, oxidative stress and HIV-1 reactivation might appear obvious, intracellular redox metabolism is dependent on many pathways, and HIV-1 reactivation is a multifactorial process, hence we expect multiple mechanisms where redox might be interlinked. In fact, our NanoString data suggest that the effect of *Mtb*-specific exosomes on HIV reactivation is likely mediated by diverse genetic factors. For example, gene expression data indicate that superoxide generation by NADPH oxidase could be one of the factors contributing to an oxidative shift in *E_GSH_*and HIV-1 reactivation upon treatment with exosomes. However, the induction of several genes encoding antioxidant enzymes contradicts the requirement of ROS for viral reactivation. It appears that cells attempt to mitigate excess ROS to ensure exosome-mediated HIV-1 reactivation without triggering global ROS-mediated cytotoxicity. Consistent with this, treatment with *Mtb*-specific exosomes induces only a modest oxidative shift in *E_GSH_* (−276 mV) of U1. Similar oxidative changes in *E_GSH_* were earlier found to uniformly reactivate HIV-1 from latency without affecting viability [20]. Low levels of ROS are known to activate HIV LTR via activation of NFκB. Agreeing to this, expression data confirm the induction of NFκBIA and SQSTM1/P62 involved in NFKB activation upon exosome treatment. The induction of AP-1 (FOS), which is activated by ROS and reactivates HIV-1 [50], also indicate that exosomes-promoted oxidative stress acts as a critical cue to reactivate HIV-1 via redox-sensitive transcription factors. This explanation aligns well with the ability of the antioxidant, NAC, in subverting HIV-1 reactivation by exosomes.

Recently, several studies have examined the contribution of metabolic indicators (OXPHOS and glycolysis) in determining susceptibility to HIV-1 infection and replication [57, 62]. Overall, these studies revealed the requirements of high OXPHOS and glycolysis rates for HIV-1 infection and continued replication [57]. In contrast, quantification of respiratory and redox parameters during HIV-1 reactivation is lacking. In this context, our data indicate a critical role of oxidative stress in reactivating HIV-1 by *Mtb* exosomes. Consistent with this, a recent study demonstrated that exosomes from *Mtb* infected neutrophils trigger superoxide production in macrophages [76]. However, the mechanism of superoxide generation remains uncharacterized. Since the respiratory chain is the major site for generation of ROS such as superoxide, cell flux assays showing *Mtb* exosome-mediated deceleration of mitochondrial OCR provide new mechanistic insight. It is well known that reduced mitochondrial OCR leads to build up of NADH, which results in trapping of flavin mononucleotide (FMN) in the reduced state on complex I of the electron transport chain [77]. Reduced FMN has been consistently shown to donate one electron to O_2_ resulting in the generation of superoxide by complex I [77]. Other sites in mitochondria, which communicate with NADH and coenzyme Q (COQ) pools such as complex III and α-ketoglutarate dehydrogenase, also produce superoxide and H_2_O_2_ during slow-down of OCR [77].

We discovered that contrary to mitochondrial OCR, non-mitochondrial OCR is significantly induced by *Mtb*-specific exosomes, which links oxidative stress with the enzymatic activities unrelated to mitochondria (e.g., NADPH oxidase, lipoxygenase, and cyclo-oxygenase) as ROS sources [58]. All of these enzymes are influenced by HIV infection and are well known to generate ROS and influence GSH homeostasis [78-80]. Importantly, a recent study showed that the treatment of bone marrow-derived macrophages (BMDM) with exosomes derived from *Mtb* infected macrophages increases recruitment of NADH oxidase on the phagosomes [60]. Taken together, data suggest that both mitochondrial and non-mitochondrial mechanisms associated with oxygen consumption likely mediate ROS generation and HIV-1 reactivation by *Mtb* exosomes.

Exosomes isolated from *Mtb* infected macrophages contain bacterial lipids, proteins, and RNA, which stimulate a pro-inflammatory response in bystander macrophages. Our study updated this knowledge by including the potential of *Mtb* exosomes in modulating redox and bioenergetics of uninfected bystander cells, which affects HIV-1 latency and reactivation program. Rather than mycobacterial components, which are well characterized, we focused on identifying the macrophage proteins enriched in *Mtb* exosomes to understand the contribution of the host on virus reactivation. Various proteins associated with host pathways mediating HIV-1 reactivation were enriched in *Mtb* exosomes. Secretion of HSP90 along with its co-chaperone Cdc37 and galectins were found to be important for *Mtb* exosomes mediated virus reactivation. HSP90 is abundantly present in the serum of HIV-TB co-infected patients [81]. Galectins such as Gal3 secretes in exosomes [82] and promotes redox imbalance [83], whereas Gal9 is frequently found in the plasma of TB patients [84] and reactivates HIV-1 [71]. While we have characterized the proteome of *Mtb*-exosomes derived from macrophages, it is certain that the presence of immuno-modulatory proteins, lipids, and RNA of *Mtb* will also influence HIV-1 reactivation.

Our findings have therapeutic implications. For example, HSP90 inhibitor, SNX-5422, shows a good safety profile in patients with solid tumors [85]. One can envisage using these inhibitors along with HAART to repress HIV-1 reactivation and replication in HIV-TB co-infected patients, at least in early stages of post-infection, when the viral reservoir is small. Furthermore, reactivation of latent virus coupled with HAART has been put forward as a possible “Kick-and-Kill” approach to eliminate latent reservoir. However, most of the screening efforts identified latency-reversing agents that are cytotoxic. Since the *Mtb*-specific exosomes mediate HIV-1 reactivation without causing overwhelming oxidative stress, we anticipate that co-treatment of *Mtb* exosomes with HAART can target HIV-1 reservoir without triggering global cytotoxicity. Exosomes derived from *Mtb* infected macrophages were already reported to potentiate the antimycobacterial activity of anti-TB drugs *in vivo* [60], suggesting that a combination of *Mtb* exosomes with HAART and/or anti-TB drugs can be exploited to reduce the burden of HIV-TB co-infection.

In conclusion, we identified new paradigms in HIV-TB co-infection, including the role of *Mtb* exosomes in inducing oxidative stress and decelerating OXHPOS in the cells latently infected with HIV-1. These events act as a signal to trigger a transcriptional response that promotes HIV-1 reactivation and inflammation. Identification of host factors enriched in *Mtb* exosomes to mediate virus reactivation has direct implications on the mechanism of HIV-1 reactivation and multiplication in HIV-TB co-infected individuals. Thus, the quantifiable redox and respiratory parameters along with the transcript and protein signatures established in this study would enable direct assessment of future antimicrobials against HIV and TB coinfection.

## Materials and methods

### Cell lines, Bacterial cultures

The human monocytic cell line U937, murine macrophage cell line RAW264.7, chronically HIV-1 infected U1 (monocytic cell line) and J1.1 (T-lymphocytic cell line) were cultured as described earlier [20, 86]. J-Lat 10.6 cells (Jurkat T-lymphocytic cell line, is a reporter cell line, containing a full-length integrated HIV-1 genome with a non-functional *env* due to frame shift and *gfp* in place of *nef* gene) were maintained in RPMI-1640 (Cell Clone) supplemented with 10% heat inactivated Fetal Bovine Serum (Sigma Aldrich), 2 mM L-glutamine, 100 U/ml penicillin and 100 mg/ml streptomycin (Sigma Aldrich) at 37°C and 5% CO2. Bacterial strains used in this study are wild type *Mycobacterium tuberculosis* H37Rv (*Mtb)*, *Mycobacterium bovis* Bacillus

Calmette Guerin (BCG), single drug resistant (SDR) clinical isolate BND 320, multiple drug resistant (MDR) clinical isolates Jal 1934, Jal 2261, and extensively drug resistant (XDR) clinical isolate MYC 431 (kind gift from Dr. Kanury V.S. Rao, ICGEB, New Delhi), were grown till mid log phase (OD_600_ of 0.8) as described previously [86]. For tdTomato expressing *Mtb* strains, competent cells were prepared as described in [86] and were electroporated using 1 μg of the pTEC27 plasmid (pMSP12::tdTomato, kind gift from Prof. Deepak Saini, IISc, Bangalore) with settings of 2.5 kV voltage, 25 μF capacitance and 1000 Ω resistance in Bio Rad Gene Pulser. Electroporated bacilli were kept for overnight recovery in 7H9 followed by selection on 7H11 agar plates containing hygromycin (50 μg/ml). After 21 days of selection, bacteria were grown in 7H9 broth till mid-log phase and used for further studies. For generating heat killed *Mtb (Hk-Mtb),* bacilli were killed by resuspending the pellet in 2 ml of RPMI and heating it to 80 °C for 30 minutes (min) using established protocol [87].

### *Mtb* labelling with PKH-26 GL, complement opsonization and infection

Freshly grown *Mtb* bacilli were labelled with fluorescent lipophilic dye PKH-26 GL (Sigma-Aldrich) as per the manufacturer’s instructions to prepare red labelled bacteria to distinguish between *Mtb* infected and bystander cells. Briefly, pelleted *Mtb* bacilli were resuspended in 300 μl of diluent C. Fluorescent staining was performed at a final concentration of 10 μM for 15 min at room temperature and fetal bovine serum (FBS) was added to terminate the labelling process. Bacilli were then washed thoroughly three times with 1X phosphate buffer saline (PBS) and resuspended in RPMI-1640 (Cell Clone). For complement opsonization *Mtb* bacilli were opsonized in 50% human serum for 30 min at 37 °C as described [88] and then washed three times in 1X PBS. For infection, U937 cells stably expressing Grx1-roGFP2 in the cytosol were seeded at a density of 0.2 million per well in 24-well plates and were differentiated into macrophages by a 24 h treatment with 5 ng/ml of phorbol 12-myristate 13-acetate (PMA; Sigma-Aldrich Co. Saint Louis, MO, USA). Cells were rested overnight following chemical differentiation to ensure that they reverted to a resting phenotype before infection. Differentiated U937 cells were then infected with *Mtb* at multiplicity of infection (moi) 10 and incubated for 4 h at 37 °C in 5% CO_2_. Cells were then treated with amikacin at 200 μg/ml for 2 h to kill extracellular bacteria. After infection cells were washed thoroughly with 1X PBS and resuspended in complete media (RPMI supplemented with 10% FCS).

### Redox potential measurements

The intracellular redox potential measurements were performed as described previously [20]. Briefly, U937 cells stably expressing Grx1-roGFP2 in the cytosol were infected with PKH-26 labelled *Mtb.* At the indicated time points, cells were treated with 10 mM N-ethylmaleimide (NEM) for 5 min and fixed with 4% paraformaldehyde (PFA) for 15 min at room temperature. After washing with 1X PBS cells were analysed using FACS Verse flow cytometer (BD Biosciences). The biosensor response was measured by analysing the ratio upon excitation at 405 and 488 nm and emission at 510 nm. Data was analysed using FACSuite software. For each experiment the minimal and maximal fluorescence ratios were determined, which correspond to 100% sensor reduction and 100% sensor oxidation, respectively. Cumene hydroperoxide (CHP, 0.5 mM) was used as the oxidant and dithiothreitol (DTT, 40 mM) as the reductant. *E_GSH_* was measured using the Nernst equation as described previously [20].

### Co-cultures of U937 macrophages with J-Lat and U1 cells

Co-cultures were performed according to earlier established protocol [89]. U937 cells were seeded at a density of 0.2 million/ml in 24 well plates and were infected with *Mtb* as described earlier. J-Lat and U1 cells were added at a density of 0.1 million/ml on uninfected or infected U937 monolayers after amikacin treatment. Co-cultures were performed in complete medium (RPMI 1640 plus 10% FCS, 2 mM L-glutamine) at 37 °C and 5% CO_2_ for 5 days (d) in case of J-Lat cells and 48 h in case of U1 cells. Fresh media was added in J-Lat cells after 48 h. At indicated time points, the supernatants containing J-Lat and U1 cells were collected and centrifuged at 1500 rpm, 5 min to harvest cells. J-Lat and U1 cells were fixed in 4% PFA for 15 minutes, centrifuged at 1500 rpm, 5 min and resuspended in 1X PBS for FACS analysis. PMA and TNF-α were used at a final concentration of 5 ng/ml and 10 ng/ml, respectively.

### Culturing of J-Lat and U1 cells in U937-conditioned media

J-Lat and U1 cells were seeded at a density of 0.1 million/ml in 24 well plates in the absence or presence of two-fold dilutions of the supernatants derived from *Mtb* uninfected or infected U937 macrophages. J-Lat and U1 cells were also cultured in two-fold dilution of the culture supernatant collected from *Mtb* infected U937 macrophages grown in presence of 10 μM of exosome secretion inhibitor (GW4869). Presence of intact *Mtb* cells in the supernatant was ruled out by passing of supernantant through 0.2 μm filter followed by confirmation of bacterial viability by plating of the supernatant. J-Lat and U1 cells were harvested at different time points and were fixed in 4% PFA for 15 minutes. After centrifugation cells were resuspended in 1X PBS for FACS analysis.

### HIV-1 p24 staining

For intracellular p24 staining, U1 cells were washed with FACS buffer (1X PBS containing 10% human serum) followed by fixation and permeabilization using the fixation/permeabilization kit (eBiosciences). Permeabilized cells were then incubated with 100 μl of 1:100 dilution of phycoerythrin-conjugated mouse anti-p24 mAb (KC57-RD1; Beckman Coulter, Inc.) for 30 min at 4 °C with intermittent mixing. After washing samples twice with FACS buffer, cells were analysed with BD FACS Verse Flow cytometer (BD Biosciences). Data was analysed using FACSuite software.

### qRT-PCR

Total RNA was isolated using RNeasy mini kit (Qiagen), according to the manufacturer’s instructions. RNA (500 ng) was reverse transcribed to cDNA using iScript^TM^ cDNA synthesis kit (Bio-Rad) using random oligonucleotide primers. p24 specific primers (p24, Forward-5′ ATAATCCACCTATCCCAGTAGGAGAAAT 3′ and Reverse-5′ TTGGTTCCTTGTCTTATGTCCAGAATGC 3′) were used to perform PCR. Gene expression was analysed with real time PCR using iQTM SYBR Green Supermix (Bio-Rad) and a CFX96 RT-PCR system (Bio-Rad). Data analysis was performed with CFXManager^TM^ software (Bio-Rad). The expression level of each gene is normalized to human β-actin (Actin, Forward-5′ATGTGGCCGAGGACTTTGATT 3′ and Reverse-5′ AGTGGGGTGGCTTTTAGGATG 3′) gene.

### Isolation and purification of exosomes

Exosomes were isolated and purified using ultrafiltration and exosome precipitation technique as described previously [90, 91]. Briefly, ∼60 million RAW264.7 macrophages were cultured (6 million cells per 100 mm cell culture dish) in cell culture dishes as described before [86] followed by infection at moi 10 with *Mtb,* Hk-*Mtb,* Jal 1934 (MDR), MYC 431 (XDR) or left uninfected. After infection, cells were washed thoroughly with 1X PBS and incubated in serum free DMEM (Cell Clone) at 37 °C and 5% CO_2_. Serum free media was used cells to avoid contamination of exosomes present in the FBS exosomes in the exosomes purified from macrophages infected with *Mtb*. Culture supernatants were collected 24 h post infection (p.i.) and were centrifuged at 5000 rpm for 15 min at 4 ℃ to remove cells, cell debris or any *Mtb* in supernatant. Cleared culture supernatants were filtered twice through 0.22 μm polyethersulphone (PES) filters (Jet Biofil^TM^). Exosomes are 30-100 nm in diameter and filter freely through 0.22 μm filters. Filtered supernatants were concentrated to 1 ml using an Amicon Ultra-15 (Merck) with a 100 kDa molecular weight cut off (MWCO) in a swing-out rotor (Thermo scientific^TM^ SL 16) at 4 ℃ and 4000 × g. An equal volume of ExoQuick (Systems BioSciences Inc. CA), exosome precipitation solution, was added to concentrated culture supernatant, and the resulting solution was mixed by inverting the tube and allowing it to stand overnight at 4 ℃. This mixture was then centrifuged at 1500 × g for 30 min. The supernatant was discarded, and the precipitate consisted of exosomes was then re-suspended in sterile 1 × PBS (filtered through 0.22 μm filter) mixed with protease inhibitor cocktail (Pierce^TM^ Thermo Fisher Scientific) and were stored at −80 ℃ for future analysis.

### Mouse infection and isolation of exosomes from serum and lungs

6 to 8 week old BALB/c mice were infected with *Mtb* by aerosol with ∼100 bacilli per mouse or left uninfected as described previously [92]. At 20 weeks p.i., animals were sacrificed and serum and lungs were collected. Exosomes were isolated from mouse serum by precipitating in ExoQuick solution overnight as per manufacturer’s instruction. To isolate exosomes from lungs, 2 ml of tissue digestion mix (Serum free RPMI with 200 μg/ml Liberase DL [Sigma-Aldrich] and 100 μg/ml of DNase [Thermo Fisher scientific]) was added to one whole lung and transferred to C-tubes. Lungs were homogenised on gentleMACS^TM^ Dissociator (Miltenyi Biotec) using machine program m_lung_01. Samples were incubated for 30 min at 37 ℃, 70-100 rpm followed by further homogenizing lungs using machine program m_lung_02 for 22 sec. Lung homogenates were passed through 40 μm cell strainer and centrifuged at 1500 rpm for 5 min. Supernatants were collected and passed through 0.22 μm filter. Filtered supernatants were concentrated to 1 mL and exosomes were isolated using ExoQuick precipitation solution as described in above section. Exosomes were stored in −80 ℃ for future analysis.

### Western blotting

Exosome were lysed in RIPA buffer (25 mM tris-HCl pH 7.6, 150 mM NaCl, 1% NP-40, 1% sodium deoxycholate and 0.1% SDS) with protease inhibitor (Pierce^TM^ Thermo Scientific). Protein estimation was performed using micro BCA assay (Pierce^TM^ Thermo Fisher Scientific). 50 μg of protein unless specified was mixed with Laemmli buffer, heated at 95 °C for 5 min followed by chilling on ice for 5 min before loading onto SDS-PAGE gel. Western blotting was performed using LAMP2 (ab25631), Rab5b (BD-610281), Alix (CST-2171), CD63 (sc-15363), p24 (ab9071), α-Tubulin (CST-2144) as primary antibodies and goat anti-Rabbit IgG HRP (CST-7074) and horse anti-mouse IgG (CST-7076) were used as secondary antibodies.

### Transmission electron microscopy

Exosomes were fixed in 4% PFA for 10 min and 10 μl sample was mounted onto a carbon formvar coated copper grid. The samples were allowed to adsorb on grids for 10 min to form a monolayer and the remaining sample was wiped off using a clean filter paper. Grids were washed thrice with 1X PBS followed by incubation in 50 μl drop of 1% glutaraldehyde for 5 min. Grids were washed thoroughly with 1X PBS and stained with 2 μl of filtered 2% uranyl acetate solution for 1 min. After washing thrice with 1X PBS grids were dried at room temperature.

For immunogold labelling of exosomes with anti-CD63 antibody, exosomes were fixed in 4% PFA. Fixed exosome samples were mounted on carbon formvar coated 300 mesh copper grids for 10 min before wiping excess using filter paper. Grids were blocked in 0.5% bovine serum albumin (BSA) in 1X PBS (blocking buffer) for 30 min and washed thrice in 1X PBS. After this, grids were incubated in blocking buffer (negative control) or primary antibody (CD63) diluted to 1:100 for 1 h. Grids were washed thoroughly with 1X PBS followed by incubation with anti-rabbit 10 nm gold antibody (ab27234) diluted to 1:250 for 1 h. Grids were then incubated in 1% glutaraldehyde for 5 min to fix the immunoreaction. Negative staining was performed using 2% aqueous uranyl acetate solution for 1 min. After washing grids were air dried and viewed with JEM 1011 transmission electron microscope at 120kV.

### NanoString gene array

U1 cells were treated with 100 μg/ml concentration of exosomes for 12 h and total RNA was isolated using RNeasy mini kit (Qiagen) according to the manufacturer’s instructions. RNA concentration and purity were measured using a Nanodrop Spectrophotometer (Thermo Fisher Scientific, Waltham, MA). nCounter Gene Expression Assay was performed according to the manufacturer’s protocol. The assay utilized a custom made NanoString codeset designed to measure 185 transcripts which includes 6 putative housekeeping transcripts (see table S1a). This custom-made panel includes genes reported to change expression in response to HIV infection and oxidative stress. The data was normalized to the average counts for all housekeeping genes in each sample and analysed in nSolver software (NanoString Technologies).

### OCR measurement

Oxygen consumption rate (OCR) was measured at 37 °C using Seahorse XFp extracellular flux analyser (Seahorse Bioscience). XF cell culture microplate plates were coated with 10 μl of Cell-Tak (Sigma-Aldrich) reagent according to the manufacturer’s protocol. U1 cells were treated with exosomes isolated from *Mtb,* Hk-*mtb* infected or uninfected RAW264.7 macrophages at 100 μg/ml concentration for 48 h. After 48 h U1 cells were washed and seeded in Seahorse flux analyser microplate pre-coated with Cell-Tak at density of 50000 cells per well to generate a confluent monolayer of cells. Agilent seahorse XFp cell mito stress kit (Agilent Technologies) was utilized to carry out mitochondrial respiration assay. Briefly, three OCR measurements were performed in XF assay media without addition of any inhibitor to measure basal respiration, followed by sequential exposure of cells to oligomycin (1 μM), an ATP synthase inhibitor and three OCR measurements to determine the ATP-linked OCR and proton leak. Then, cyanide-4-[trifluoromethoxy]phenylhydrazone (FCCP; 0.25 μM), an Electron transport chain (ETC) uncoupler was injected to determine the maximal respiration and the spare respiratory capacity (SRC). Lastly, antimycin A and rotenone (0.5 μM each) inhibitor of complex III and I; respectively, were injected to completely shut down the ETC to determine non-mitochondrial respiration. The Wave Desktop 2.6 Software from Agilent website (https://www.agilent.com/en/products/cell-analysis/software-download-for-wave-desktop) was used for the calculation of the parameters from mitochondrial respiration assay. Data was normalized according to protocol described previously [93].

### Proteomic analysis of exosomes by LC-MS/MS

We extracted proteins from exosomes as described previously for the immuno-blotting experiment. Protein samples (30 μg) were resolved on 10% SDS-PAGE gel up to a distance of 3 cm and stained with Coomassie Brilliant Blue R250. The lanes were cut into three equal size bands. These bands were first reduced with 5 mM Tris (2-carboxyethyl) phosphine hydrochloride (TCEP; Sigma-Aldrich) followed by alkylation with 50 mM iodoacetamide and digested with 1μg trypsin for as long as 16 hours at 37 °C. The digests were then cleaned up using C18 silica cartridge (The Nest Group, Southborough, MA) and dried using speed vac which then was resuspended in Buffer A (5% acetonitrile / 0.1% formic acid).

EASY-nLC 1000 system (Thermo Fisher Scientific) was used to perform LC-MS/MS, coupled to QExactive mass spectrometer (Thermo Fisher Scientific) fitted with nanoelectrospray ion source. 15 cm Pico-Frit filled with 1.8 μm of C18 resin (Dr. Maeisch) was used to load and resolve 1 μg of the peptide mixture with Buffer A. Loading and elution with 0-40% gradient of Buffer B (95% acetonitrile/0.1% Formic acid) was given a flow rate of 300 nl/min and RT of 105 minutes. The MS was driven with a full scan resolution of 70,000 at m/z of 400 and the MS/MS scans were acquired at a resolution of 17,500 at mz of 400 using Top10 HCD Data-dependant acquisition mode. Polydimethylcyclosiloxane (PCM) ions (m/z = 445.120025) was set up as lock mass option for internal recaliberation during the run. MS Data acquisition was carried out using a Data-dependent Top10 method, which effectively chooses most abundant precursor ions from a survey scan.

Raw files were analyzed using Thermo Proteome Dicoverer 2.2 searched against Uniprot Mus musculus reference proteome database with both PSM (peptide spectrum matches) and protein FDR set to 0.01 using percolator node. For Sequest HT search, the precursor and fragment mass tolerances were set at 10 ppm and 0.5 Da, respectively. Protein quantification was done using Minora feature detector node with default settings and considering only high PSM (peptide spectrum matches) confidence.

### Data processing and analysis

Differential analysis was performed on Label-free quantification data using R packages-ProstaR and DAPAR. The intensity values were log-transformed followed by filtering of rows containing *NA*>=5 in the data. Imputation was performed using R package-Mice. Limma-moderated t-test was used to identify differentially expressed proteins and p values were adjusted using BH method, between two groups. All proteins with fold change >2 or <−2 and with *p*value<0.01 were considered significant. Functional classification of differentially expressed proteins with GO and KEGG signalling pathway between two groups were analyzed using R package-clusterProfiler.

### Statistical analysis

Statistical analyses were performed using the GraphPad Prism software. Statistical analyses were performed using Student’s *t*-tests (two-tailed). Comparisons of multiple groups were made by either using one-way or two-way ANOVA with Bonferroni multiple comparisons. Differences with a *p* value of < 0.05 were considered significant.

### Ethics statement

This study was carried out in strict accordance with the guidelines provided by the Committee for the Purpose of Control and Supervision on Experiments on Animals (CPCSEA), Government of India. The protocol was approved by the Animal Ethics Committee (AEC) of Indian Institute of Science (#CAF/Ethics/485/2016).

## Acknowledgments

This work was supported by the Wellcome trust-DBT India Alliance grant IA/S/16/2/502700 (AS), and in part by Department of Biotechnology (DBT) Grant BT/PR11911/BRB/10/1327/2014, BT/PR13522/COE/34/27/2015 (AS), DBT-IISc Partnership Program (22-0905-0006-05-987 436) and Infosys foundation. AS is a senior fellow of Wellcome trust-DBT India Alliance. PT and VKP acknowledge fellowships from CSIR and IISc, respectively.

## Supplementary Information

### Supplementary figures

**Fig. S1.**
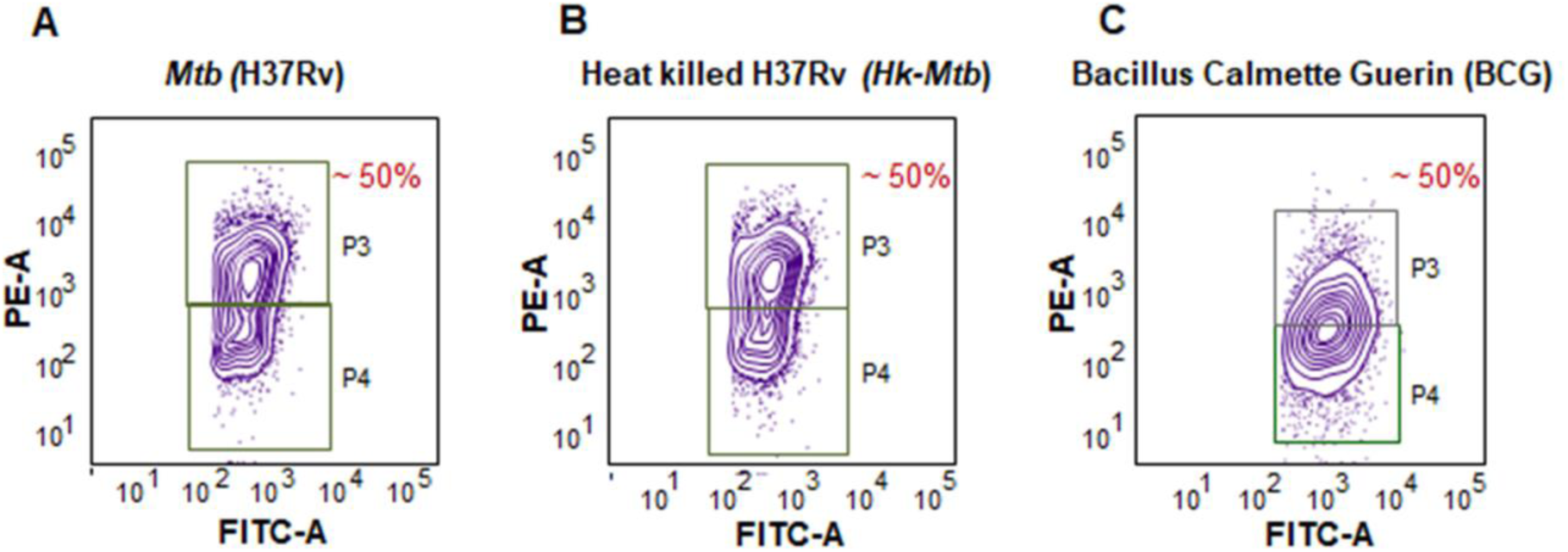
*Mtb* H37Rv, Heat killed H37Rv (*Hk-Mtb)*, and BCG equally infect the U937 Mφ. PMA differentiated U937/Grx1-roGFP2 were infected with PKH26 labeled bacilli (MOI 10). **(A-C)** Dot plot of U937/Grx1-roGFP2 after infection with **(A)** H37Rv (*Mtb),* **(B)** (Hk-*Mtb*) and **(C)** Bacillus Calmette Guerin (BCG). P3 represents *Mtb* infected U937 cell subpopulation and P4 represent bystander U937 cell subpopulation.

**Fig. S2.**
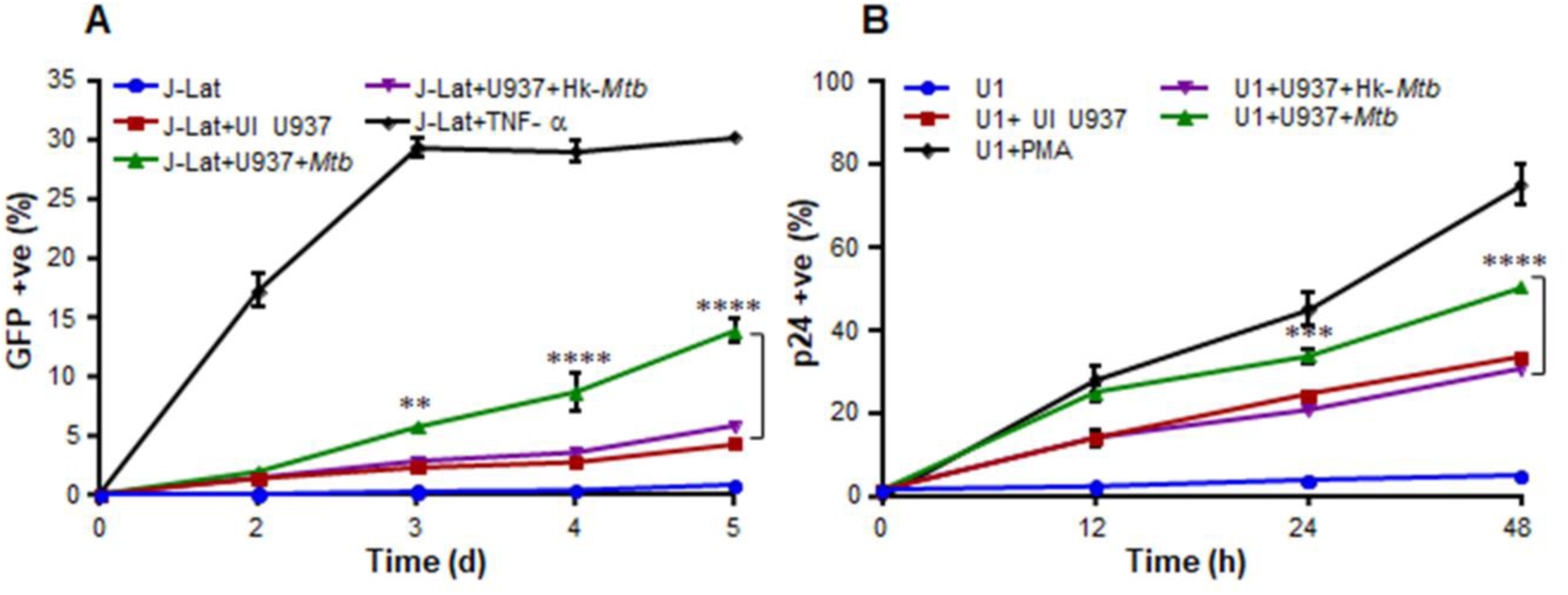
*Mtb* infected Mφ reactivates latent HIV-1 from lymphocytes and monocytes. **(A)** J-Lat T cells are latently infected with the HIV-1 provirus containing green fluorescent protein [GFP] in the genome. Reactivation of HIV-1 can be easily monitored by measuring the increase in GFP fluorescence using flow cytometer. J-Lat cells were co-cultured with U937 Mφ pre-infected with live *Mtb* or *Hk*-*Mtb* for 4 h and GFP fluorescence was monitored at the indicated time points. GFP fluorescence of J-Lat cells treated with TNFα (10 ng/ml) or left untreated was taken as the measure of reactivation or latency, respectively. As an additional control, J-Lat cells were also co-cultured with uninfected (UI) U937 Mφ. **(B)** U1 cells (U937 monocytes chronically infected with HIV-1), were co-cultured with UI, *Mtb* infected or Hk-*Mtb* infected U937 Mφ. At various time points, U1 cells were immuno-stained for intracellular HIV p24 (Gag) antigen and fluorescence response was measured by flow cytometry. Percent p24 antigen positive U1 cells corresponds to percent HIV reactivation upon co-culturing over time. PMA (5 ng/ml) and U1 cells alone were used as positive and negative control, respectively. Error bars represent standard deviations from the mean. ** p<0.01; ***p<0.001; **** p<0.0001 (two-way ANOVA). Data are representative of at least three independent experiments performed in triplicate.

**Fig. S3.**
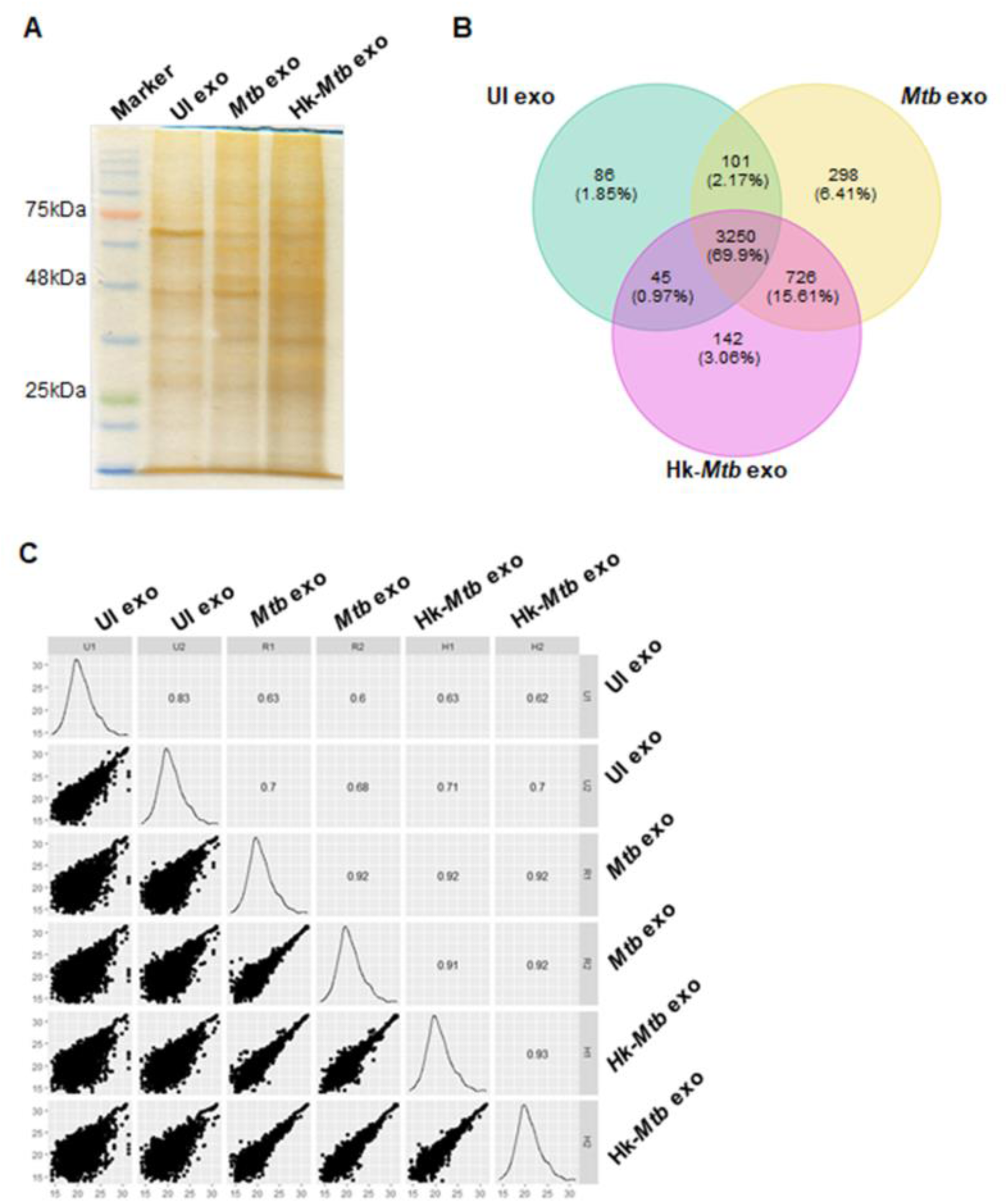
Proteomics of *Mtb* specific exosomes by LC-MS/MS. RAW264.7 Mφ were infected with *Mtb* and Hk-*Mtb* at moi 10 or left uninfected. At 24 h p.i., exosomes were isolated from the culture supernatant for LC-MS/MS. **(A)** Silver stained SDS-PAGE gel loaded with 10 μg of total exosomal protein. **(B)** Venn diagram showing percent overlap between the exosomal proteins identified by LC-MS/MS among three samples. **(C)** Correlation plot of three exosome samples in duplicate.

**Fig. S4.**
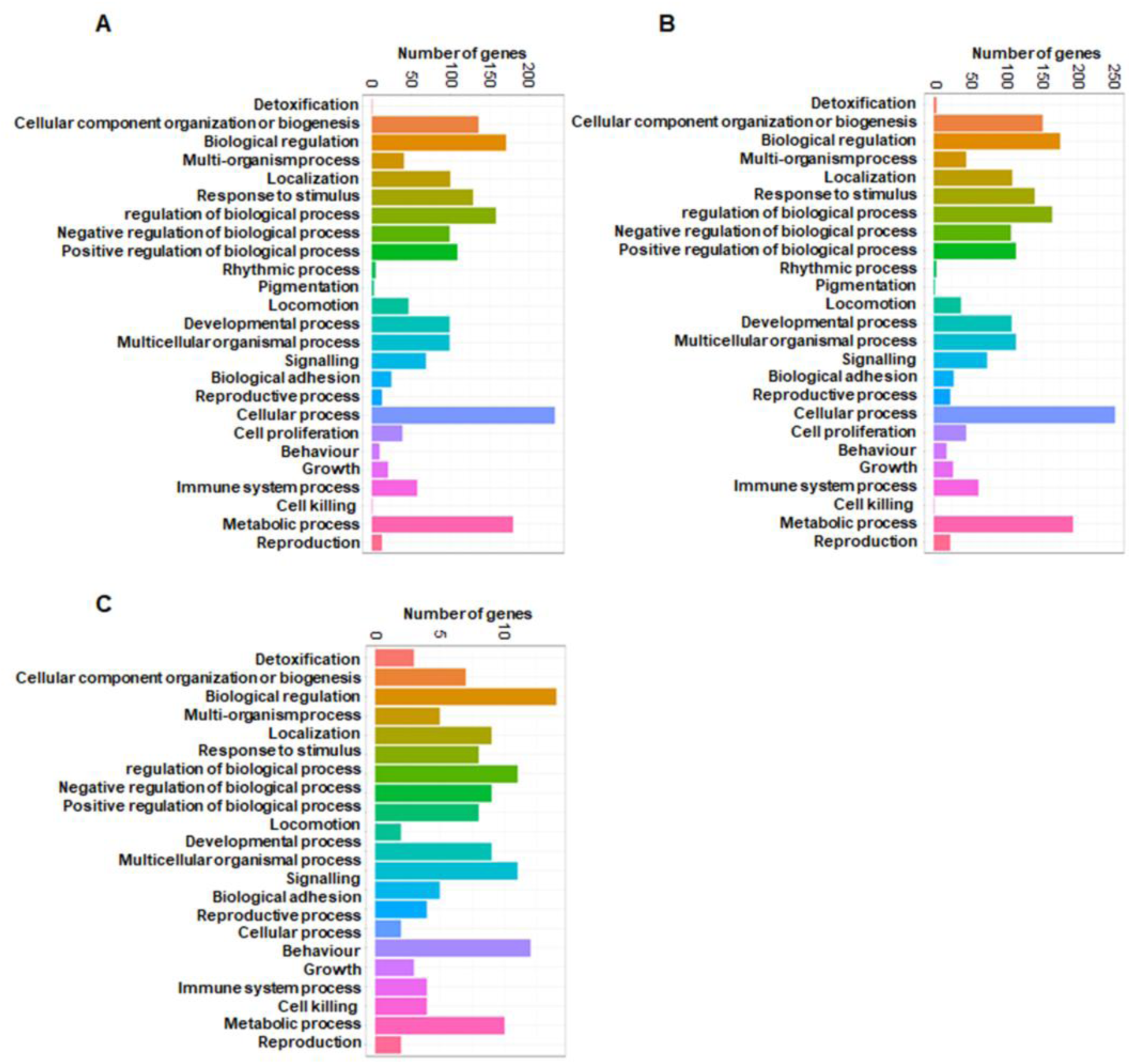
Functional classification of differentially expressed proteins of exosomes. **(A)** Enriched GO terms (biological processes) of differentially expressed proteins in *Mtb* exosomes *versus* UI exosomes. **(B)** Enriched GO terms (biological processes) of differentially expressed proteins in Hk-*Mtb* exosomes *versus* UI exosomes. **(C)** Enriched GO terms (biological processes) of differentially expressed proteins in *Mtb* exosomes *versus* Hk-*Mtb* exosomes.

**Fig. S5.**
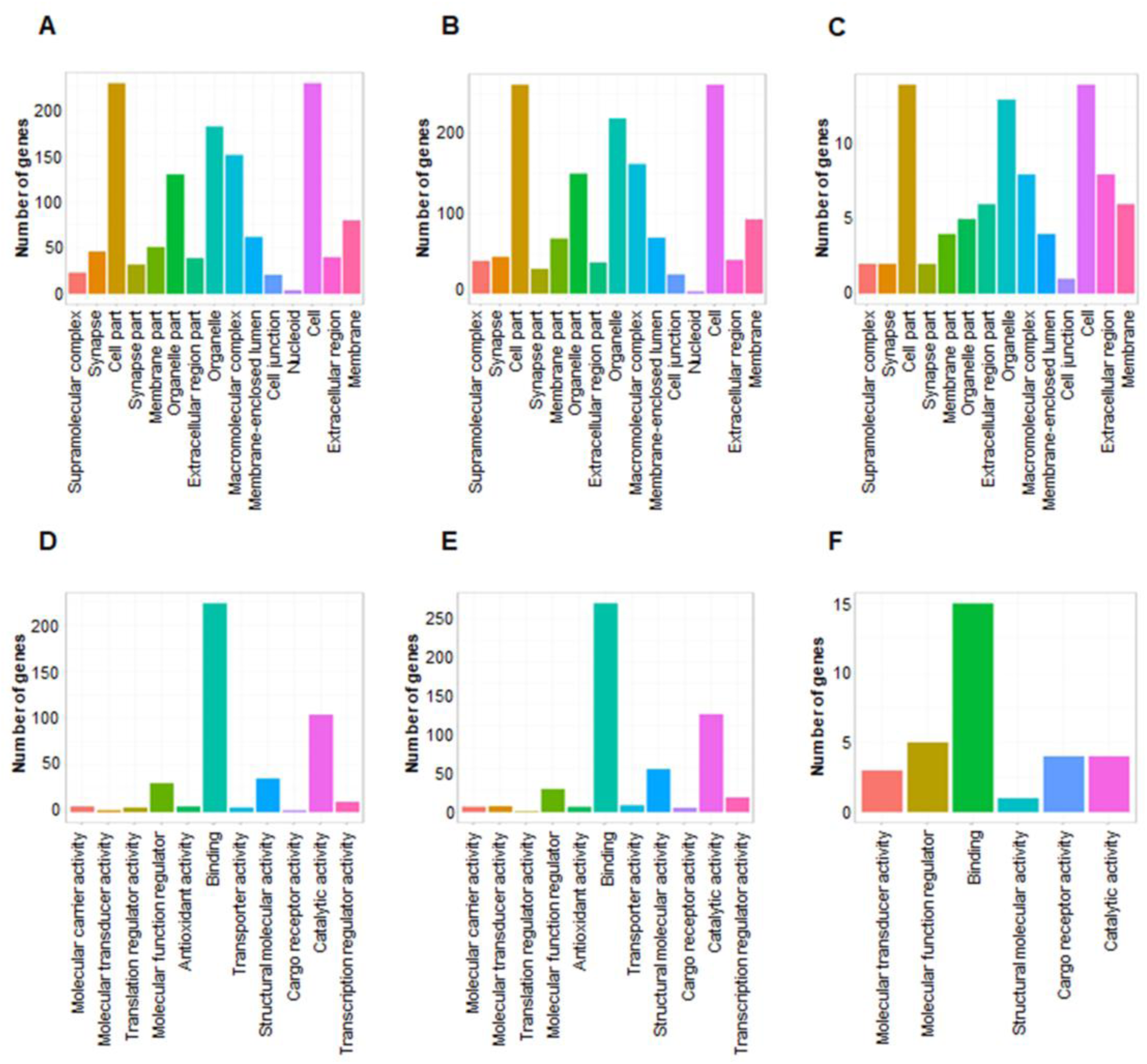
Functional classification of differentially expressed proteins of exosomes. **(A)** Enriched GO terms (Cellular localization) of differentially expressed proteins in *Mtb* exosomes *versus* UI exosomes. **(B)** Enriched GO terms (Cellular localization) of differentially expressed proteins in Hk-*Mtb* exosomes *versus* UI exosomes. **(C)** Enriched GO terms (Cellular localization) of differentially expressed proteins in *Mtb* exosomes *versus* Hk-*Mtb* exosomes. **(D)** Enriched GO terms (Molecular Function) of differentially expressed proteins in *Mtb* exosomes *versus* UI exosomes. **(E)** Enriched GO terms (Molecular Function) of differentially expressed proteins in Hk-*Mtb* exosomes *versus* UI exosomes. **(F)** Enriched GO terms (Molecular function) of differentially expressed proteins in *Mtb* exosomes *versus* Hk-*Mtb* exosomes.

## Tables

### Table S1

**Table S1a.** List of genes used for nCounter Gene Expression Assay.

**Table S1b.** Normalized intensity values of differentially expressed gene in nCounter Gene Expression Assay.

**Table S1c.** Normalized intensity values of differentially expressed gene in nCounter Gene Expression Assay.

**Table S2.** List of proteins identified in LC/MS-MS of uninfected, live H37Rv (*Mtb*) and heat killed H37Rv (*Hk-Mtb)* exosomes.

### Table S3

**Table S3a.** List of differentially expressed proteins in live H37Rv *(Mtb)* versus Uninfected (UI) exosomes shown in volcano plots.

**Table S3b.** List of differentially expressed proteins in heat killed H37Rv *(Hk-Mtb)* versus Uninfected (UI) exosomes shown in volcano plots.

**Table S3c.** List of differentially expressed proteins in heat killed H37Rv *(Hk-Mtb)* versus live H37Rv (*Mtb*) exosomes shown in volcano plots.

### Table S4

**Table S4a.** Biological process Gene ontology (GO) term enrichment analysis of live *H37Rv (Mtb)* versus Uninfected (UI) exosome proteins.

**Table S4b.** Cell localisation Gene ontology (GO) term enrichment analysis of live *H37Rv (Mtb)* versus Uninfected (UI) exosome proteins.

**Table S4c.** Molecular function Gene ontology (GO) term enrichment analysis of live *H37Rv (Mtb)* versus Uninfected (UI) exosome proteins.

### Table S5

**Table S5a.** Biological process Gene ontology (GO) term enrichment analysis of heat killed *H37Rv (Hk-Mtb)* versus Uninfected (UI) exosome proteins.

**Table S5b.** Cell localisation Gene ontology (GO) term enrichment analysis of heat killed H37Rv (Hk-Mtb) versus Uninfected (UI) exosome proteins.

**Table S5c.** Molecular function Gene ontology (GO) term enrichment analysis of heat killed H37Rv (Hk-Mtb) versus Uninfected (UI) exosome proteins.

### Table 6

**Table S6a.** Biological process Gene ontology (GO) term enrichment analysis of *live H37Rv (Mtb)* versus Heat killed H37Rv (Hk-Mtb) exosome proteins.

**Table S6b.** Cell localisation Gene ontology (GO) term enrichment analysis of *live H37Rv (Mtb)* versus Heat killed H37Rv (Hk-Mtb) exosome proteins.

**Table S6c.** Molecular function Gene ontology (GO) term enrichment analysis of *live H37Rv (Mtb)* versus Heat killed H37Rv (Hk-Mtb) exosome proteins.

**Table S7.** KEGG pathway enrichment analysis of heat killed H37Rv (H) *exosomes* versus live H37Rv (R) exosomes, heat killed H37Rv (H) exosomes versus Uninfected (U) exosomes, live H37Rv (R) versus Uninfected (U) exosome proteins.

## References

1. Whalen, C., et al., Accelerated course of human immunodeficiency virus infection after tuberculosis. Am J Respir Crit Care Med, 1995. 151(1): p. 129–35.

2. Kwan, C.K. and J.D. Ernst, HIV and tuberculosis: a deadly human syndemic. Clin Microbiol Rev, 2011. 24(2): p. 351–76.

3. DeRiemer, K., et al., Quantitative impact of human immunodeficiency virus infection on tuberculosis dynamics. Am J Respir Crit Care Med, 2007. 176(9): p. 936–44.

4. Nusbaum, R.J., et al., Pulmonary Tuberculosis in Humanized Mice Infected with HIV-1. Sci Rep, 2016. 6: p. 21522.

5. Mwandumba, H.C., et al., Mycobacterium tuberculosis resides in nonacidified vacuoles in endocytically competent alveolar macrophages from patients with tuberculosis and HIV infection. J Immunol, 2004. 172(7): p. 4592–8.

6. Kalsdorf, B., et al., HIV-1 infection impairs the bronchoalveolar T-cell response to mycobacteria. Am J Respir Crit Care Med, 2009. 180(12): p. 1262–70.

7. Diedrich, C.R. and J.L. Flynn, HIV-1/mycobacterium tuberculosis coinfection immunology: how does HIV-1 exacerbate tuberculosis? Infect Immun, 2011. 79(4): p. 1407–17.

8. Bafica, A., et al., Cutting edge: in vivo induction of integrated HIV-1 expression by mycobacteria is critically dependent on Toll-like receptor 2. J Immunol, 2003. 171(3): p. 1123–7.

9. Toossi, Z., et al., Increased replication of HIV-1 at sites of Mycobacterium tuberculosis infection: potential mechanisms of viral activation. J Acquir Immune Defic Syndr, 2001. 28(1): p. 1–8.

10. Lederman, M.M., et al., Mycobacterium tuberculosis and its purified protein derivative activate expression of the human immunodeficiency virus. J Acquir Immune Defic Syndr, 1994. 7(7): p. 727–33.

11. Zhang, Y., et al., Mycobacterium tuberculosis enhances human immunodeficiency virus-1 replication by transcriptional activation at the long terminal repeat. J Clin Invest, 1995. 95(5): p. 2324–31.

12. Bernier, R., et al., Mycobacterium tuberculosis mannose-capped lipoarabinomannan can induce NF-kappaB-dependent activation of human immunodeficiency virus type 1 long terminal repeat in T cells. J Gen Virol, 1998. 79 (Pt 6): p. 1353–61.

13. Souriant, S., et al., Tuberculosis Exacerbates HIV-1 Infection through IL-10/STAT3-Dependent Tunneling Nanotube Formation in Macrophages. Cell Rep, 2019. 26(13): p. 3586–3599 e7.

14. Goletti, D., et al., Inhibition of HIV-1 replication in monocyte-derived macrophages by Mycobacterium tuberculosis. J Infect Dis, 2004. 189(4): p. 624–33.

15. Ranjbar, S., et al., HIV-1 replication is differentially regulated by distinct clinical strains of Mycobacterium tuberculosis. PLoS One, 2009. 4(7): p. e6116.

16. Perl, A. and K. Banki, Genetic and metabolic control of the mitochondrial transmembrane potential and reactive oxygen intermediate production in HIV disease. Antioxid Redox Signal, 2000. 2(3): p. 551–73.

17. Herzenberg, L.A., et al., Glutathione deficiency is associated with impaired survival in HIV disease. Proc Natl Acad Sci U S A, 1997. 94(5): p. 1967–72.

18. Peterson, J.D., et al., Glutathione levels in antigen-presenting cells modulate Th1 versus Th2 response patterns. Proc Natl Acad Sci U S A, 1998. 95(6): p. 3071–6.

19. De Rosa, S.C., et al., N-acetylcysteine replenishes glutathione in HIV infection. Eur J Clin Invest, 2000. 30(10): p. 915–29.

20. Bhaskar, A., et al., Measuring glutathione redox potential of HIV-1-infected macrophages. J Biol Chem, 2015. 290(2): p. 1020–38.

21. Guerra, C., et al., Glutathione and adaptive immune responses against Mycobacterium tuberculosis infection in healthy and HIV infected individuals. PLoS One, 2011. 6(12): p. e28378.

22. Palanisamy, G.S., et al., Evidence for oxidative stress and defective antioxidant response in guinea pigs with tuberculosis. PLoS One, 2011. 6(10): p. e26254.

23. Cumming, B.M., et al., Mycobacterium tuberculosis induces decelerated bioenergetic metabolism in human macrophages. Elife, 2018. 7.

24. Gutscher, M., et al., Real-time imaging of the intracellular glutathione redox potential. Nat Methods, 2008. 5(6): p. 553–9.

25. Choi, H.H., et al., Endoplasmic reticulum stress response is involved in Mycobacterium tuberculosis protein ESAT-6-mediated apoptosis. FEBS Lett, 2010. 584(11): p. 2445–54.

26. Shenoi, S., et al., Multidrug-resistant and extensively drug-resistant tuberculosis: consequences for the global HIV community. Curr Opin Infect Dis, 2009. 22(1): p. 11–7.

27. Wells, C.D., et al., HIV infection and multidrug-resistant tuberculosis: the perfect storm. J Infect Dis, 2007. 196 Suppl 1: p. S86–107.

28. Kumar, D., et al., Genome-wide analysis of the host intracellular network that regulates survival of Mycobacterium tuberculosis. Cell, 2010. 140(5): p. 731–43.

29. Jordan, A., D. Bisgrove, and E. Verdin, HIV reproducibly establishes a latent infection after acute infection of T cells in vitro. EMBO J, 2003. 22(8): p. 1868–77.

30. Folks, T.M., et al., Cytokine-induced expression of HIV-1 in a chronically infected promonocyte cell line. Science, 1987. 238(4828): p. 800–2.

31. Poli, G., et al., Tumor necrosis factor alpha functions in an autocrine manner in the induction of human immunodeficiency virus expression. Proc Natl Acad Sci U S A, 1990. 87(2): p. 782–5.

32. Kurosaka, K., N. Watanabe, and Y. Kobayashi, Production of proinflammatory cytokines by phorbol myristate acetate-treated THP-1 cells and monocyte-derived macrophages after phagocytosis of apoptotic CTLL-2 cells. J Immunol, 1998. 161(11): p. 6245–9.

33. Bhatnagar, S., et al., Exosomes released from macrophages infected with intracellular pathogens stimulate a proinflammatory response in vitro and in vivo. Blood, 2007. 110(9): p. 3234–44.

34. Chan, J.K. and W.C. Greene, NF-kappaB/Rel: agonist and antagonist roles in HIV-1 latency. Curr Opin HIV AIDS, 2011. 6(1): p. 12–8.

35. Essandoh, K., et al., Blockade of exosome generation with GW4869 dampens the sepsis-induced inflammation and cardiac dysfunction. Biochim Biophys Acta, 2015. 1852(11): p. 2362–71.

36. Perez, V.L., et al., An HIV-1-infected T cell clone defective in IL-2 production and Ca2+ mobilization after CD3 stimulation. J Immunol, 1991. 147(9): p. 3145–8.

37. Singh, P.P., et al., Exosomes isolated from mycobacteria-infected mice or cultured macrophages can recruit and activate immune cells in vitro and in vivo. J Immunol, 2012. 189(2): p. 777–85.

38. Booth, A.M., et al., Exosomes and HIV Gag bud from endosome-like domains of the T cell plasma membrane. J Cell Biol, 2006. 172(6): p. 923–35.

39. Lee, Y., S. El Andaloussi, and M.J. Wood, Exosomes and microvesicles: extracellular vesicles for genetic information transfer and gene therapy. Hum Mol Genet, 2012. 21(R1): p. R125–34.

40. Staal, F.J., et al., Intracellular thiols regulate activation of nuclear factor kappa B and transcription of human immunodeficiency virus. Proc Natl Acad Sci U S A, 1990. 87(24): p. 9943–7.

41. Pyo, C.W., et al., Reactive oxygen species activate HIV long terminal repeat via post-translational control of NF-kappaB. Biochem Biophys Res Commun, 2008. 376(1): p. 180–5.

42. Kulkarni, M.M., Digital multiplexed gene expression analysis using the NanoString nCounter system. Curr Protoc Mol Biol, 2011. Chapter 25: p. Unit25B 10.

43. Salzano, S., et al., Linkage of inflammation and oxidative stress via release of glutathionylated peroxiredoxin-2, which acts as a danger signal. Proc Natl Acad Sci U S A, 2014. 111(33): p. 12157–62.

44. Volonte, D. and F. Galbiati, Inhibition of thioredoxin reductase 1 by caveolin 1 promotes stress-induced premature senescence. EMBO Rep, 2009. 10(12): p. 1334–40.

45. Yang, X., et al., Mn Inhibits GSH Synthesis via Downregulation of Neuronal EAAC1 and Astrocytic xCT to Cause Oxidative Damage in the Striatum of Mice. Oxid Med Cell Longev, 2018. 2018: p. 4235695.

46. Lee, J.H., et al., Ferritin binds and activates p53 under oxidative stress. Biochem Biophys Res Commun, 2009. 389(3): p. 399–404.

47. Gill, A.J., et al., Heme oxygenase-1 promoter region (GT)n polymorphism associates with increased neuroimmune activation and risk for encephalitis in HIV infection. J Neuroinflammation, 2018. 15(1): p. 70.

48. Gao, Y., et al., Activation of the selenoprotein SEPS1 gene expression by pro-inflammatory cytokines in HepG2 cells. Cytokine, 2006. 33(5): p. 246–51.

49. Nakamura, K., et al., PB1 domain interaction of p62/sequestosome 1 and MEKK3 regulates NF-kappaB activation. J Biol Chem, 2010. 285(3): p. 2077–89.

50. Roebuck, K.A., D.S. Gu, and M.F. Kagnoff, Activating protein-1 cooperates with phorbol ester activation signals to increase HIV-1 expression. AIDS, 1996. 10(8): p. 819–26.

51. Mamik, M.K. and A. Ghorpade, Chemokine CXCL8 promotes HIV-1 replication in human monocyte-derived macrophages and primary microglia via nuclear factor-kappaB pathway. PLoS One, 2014. 9(3): p. e92145.

52. Ansari, A.W., A. Kamarulzaman, and R.E. Schmidt, Multifaceted Impact of Host C-C Chemokine CCL2 in the Immuno-Pathogenesis of HIV-1/M. tuberculosis Co-Infection. Front Immunol, 2013. 4: p. 312.

53. Albin, J.S. and R.S. Harris, Interactions of host APOBEC3 restriction factors with HIV-1 in vivo: implications for therapeutics. Expert Rev Mol Med, 2010. 12: p. e4.

54. Sadler, H.A., et al., APOBEC3G contributes to HIV-1 variation through sublethal mutagenesis. J Virol, 2010. 84(14): p. 7396–404.

55. Toossi, Z., et al., Short Communication: Expression of APOBEC3G and Interferon Gamma in Pleural Fluid Mononuclear Cells from HIV/TB Dual Infected Subjects. AIDS Res Hum Retroviruses, 2015. 31(7): p. 692–5.

56. Santos, S., et al., Virus-producing cells determine the host protein profiles of HIV-1 virion cores. Retrovirology, 2012. 9: p. 65.

57. Hegedus, A., M. Kavanagh Williamson, and H. Huthoff, HIV-1 pathogenicity and virion production are dependent on the metabolic phenotype of activated CD4+ T cells. Retrovirology, 2014. 11: p. 98.

58. Chacko, B.K., et al., The Bioenergetic Health Index: a new concept in mitochondrial translational research. Clin Sci (Lond), 2014. 127(6): p. 367–73.

59. Schorey, J.S., et al., Exosomes and other extracellular vesicles in host-pathogen interactions. EMBO Rep, 2015. 16(1): p. 24–43.

60. Cheng, Y. and J.S. Schorey, Extracellular vesicles deliver Mycobacterium RNA to promote host immunity and bacterial killing. EMBO Rep, 2019. 20(3).

61. Rottenberg, D.A., et al., Abnormal cerebral glucose metabolism in HIV-1 seropositive subjects with and without dementia. J Nucl Med, 1996. 37(7): p. 1133–41.

62. Palmer, C.S., et al., Glucose Metabolism in T Cells and Monocytes: New Perspectives in HIV Pathogenesis. EBioMedicine, 2016. 6: p. 31–41.

63. Townsend, D.M., K.D. Tew, and H. Tapiero, Sulfur containing amino acids and human disease. Biomed Pharmacother, 2004. 58(1): p. 47–55.

64. Rao, S., et al., Host mRNA decay proteins influence HIV-1 replication and viral gene expression in primary monocyte-derived macrophages. Retrovirology, 2019. 16(1): p. 3.

65. Kok, Y.L., et al., Spontaneous reactivation of latent HIV-1 promoters is linked to the cell cycle as revealed by a genetic-insulators-containing dual-fluorescence HIV-1-based vector. Sci Rep, 2018. 8(1): p. 10204.

66. Felty, Q., et al., Estrogen-induced mitochondrial reactive oxygen species as signal-transducing messengers. Biochemistry, 2005. 44(18): p. 6900–9.

67. Das, B., et al., Estrogen receptor-1 is a key regulator of HIV-1 latency that imparts gender-specific restrictions on the latent reservoir. Proc Natl Acad Sci U S A, 2018. 115(33): p. E7795–E7804.

68. Duette, G., et al., Induction of HIF-1alpha by HIV-1 Infection in CD4(+) T Cells Promotes Viral Replication and Drives Extracellular Vesicle-Mediated Inflammation. MBio, 2018. 9(5).

69. Anand, P.K., et al., Exosomal Hsp70 induces a pro-inflammatory response to foreign particles including mycobacteria. PLoS One, 2010. 5(4): p. e10136.

70. Anderson, I., et al., Heat shock protein 90 controls HIV-1 reactivation from latency. Proc Natl Acad Sci U S A, 2014. 111(15): p. E1528–37.

71. Abdel-Mohsen, M., et al., Human Galectin-9 Is a Potent Mediator of HIV Transcription and Reactivation. PLoS Pathog, 2016. 12(6): p. e1005677.

72. Aalinkeel, R., et al., Galectin-1 Reduces Neuroinflammation via Modulation of Nitric Oxide-Arginase Signaling in HIV-1 Transfected Microglia: a Gold Nanoparticle-Galectin-1 “Nanoplex” a Possible Neurotherapeutic? J Neuroimmune Pharmacol, 2017. 12(1): p. 133–151.

73. Chaudhuri, A., et al., STAT1 signaling modulates HIV-1-induced inflammatory responses and leukocyte transmigration across the blood-brain barrier. Blood, 2008. 111(4): p. 2062–72.

74. Tjernlund, A., et al., Leukemia inhibitor factor (LIF) inhibits HIV-1 replication via restriction of stat 3 activation. AIDS Res Hum Retroviruses, 2007. 23(3): p. 398–406.

75. Barondes, S.H., et al., Galectins: a family of animal beta-galactoside-binding lectins. Cell, 1994. 76(4): p. 597–8.

76. Alvarez-Jimenez, V.D., et al., Extracellular Vesicles Released from Mycobacterium tuberculosis-Infected Neutrophils Promote Macrophage Autophagy and Decrease Intracellular Mycobacterial Survival. Front Immunol, 2018. 9: p. 272.

77. Murphy, M.P., How mitochondria produce reactive oxygen species. Biochem J, 2009. 417(1): p. 1–13.

78. Vilhardt, F., et al., The HIV-1 Nef protein and phagocyte NADPH oxidase activation. J Biol Chem, 2002. 277(44): p. 42136–43.

79. Kim, C., J.Y. Kim, and J.H. Kim, Cytosolic phospholipase A(2), lipoxygenase metabolites, and reactive oxygen species. BMB Rep, 2008. 41(8): p. 555–9.

80. Coffey, M.J., et al., Granulocyte-macrophage colony-stimulating factor upregulates reduced 5-lipoxygenase metabolism in peripheral blood monocytes and neutrophils in acquired immunodeficiency syndrome. Blood, 1999. 94(11): p. 3897–905.

81. Chen, C., et al., Identification of a Novel Serum Biomarker for Tuberculosis Infection in Chinese HIV Patients by iTRAQ-Based Quantitative Proteomics. Front Microbiol, 2018. 9: p. 330.

82. Banfer, S., et al., Molecular mechanism to recruit galectin-3 into multivesicular bodies for polarized exosomal secretion. Proc Natl Acad Sci U S A, 2018. 115(19): p. E4396–E4405.

83. Fulton, D.J.R., et al., Galectin-3: A Harbinger of Reactive Oxygen Species, Fibrosis, and Inflammation in Pulmonary Arterial Hypertension. Antioxid Redox Signal, 2019.

84. Zhao, J., et al., Secretion of IFN-gamma Associated with Galectin-9 Production by Pleural Fluid Cells from a Patient with Extrapulmonary Tuberculosis. Int J Mol Sci, 2017. 18(7).

85. Infante, J.R., et al., Phase I dose-escalation studies of SNX-5422, an orally bioavailable heat shock protein 90 inhibitor, in patients with refractory solid tumours. Eur J Cancer, 2014. 50(17): p. 2897–904.

86. Bhaskar, A., et al., Reengineering redox sensitive GFP to measure mycothiol redox potential of Mycobacterium tuberculosis during infection. PLoS Pathog, 2014. 10(1): p. e1003902.

87. Srivastava, M., et al., Mediator responses of alveolar macrophages and kinetics of mononuclear phagocyte subset recruitment during acute primary and secondary mycobacterial infections in the lungs of mice. Cell Microbiol, 2007. 9(3): p. 738–52.

88. Malik, Z.A., G.M. Denning, and D.J. Kusner, Inhibition of Ca(2+) signaling by Mycobacterium tuberculosis is associated with reduced phagosome-lysosome fusion and increased survival within human macrophages. J Exp Med, 2000. 191(2): p. 287–302.

89. Borghi, M.O., et al., Interaction between chronically HIV-infected promonocytic cells and human umbilical vein endothelial cells: role of proinflammatory cytokines and chemokines in viral expression modulation. Clin Exp Immunol, 2000. 120(1): p. 93–100.

90. Zeringer, E., et al., Strategies for isolation of exosomes. Cold Spring Harb Protoc, 2015. 2015(4): p. 319–23.

91. Li, P., et al., Progress in Exosome Isolation Techniques. Theranostics, 2017. 7(3): p. 789–804.

92. Mishra, S., et al., Efficacy of beta-lactam/beta-lactamase inhibitor combination is linked to WhiB4-mediated changes in redox physiology of Mycobacterium tuberculosis. 2017. 6.

93. Yang, Y., et al., Metabolic reprogramming for producing energy and reducing power in fumarate hydratase null cells from hereditary leiomyomatosis renal cell carcinoma. PLoS One, 2013. 8(8): p. e72179.

